# Dynamics of replication origin over-activation

**DOI:** 10.1101/2020.01.27.922211

**Authors:** Haiqing Fu, Christophe E. Redon, Koichi Utani, Bhushan L. Thakur, Sangmin Jang, Anna B. Marks, Sophie Z. Zhuang, Sarah B. Lazar, Mishal Rao, Shira Mencer, Jacob M. Gross, Lorinc S. Pongor, Mirit I. Aladjem

**Author notes:** Correspondence should be addressed to: M. I. Aladjem.

## Abstract

We determined replication patterns in cancer cells in which the controls that normally prevent excess replication were disrupted (“re-replicating cells”). Single-fiber analyses suggested that replication origins were activated at a higher frequency in re-replicating cells. However, nascent strand sequencing demonstrated that re-replicating cells utilized the same pool of potential replication origins as normally replicating cells. Surprisingly, re-replicating cells exhibited a skewed initiation frequency correlating with replication timing. These patterns differed from the replication profiles observed in non-re-replicating cells exposed to replication stress, which activated a novel group of dormant origins not typically activated during normal mitotic growth. Hence, disruption of the molecular interactions that regulates origin initiation can activate two distinct pools of potential replication origins: re-replicating cells over-activate flexible origins while replication stress in normal mitotic growth activates dormant origins.

## Introduction

Proliferating cells ensure the accurate transmission of their genetic material to daughter cells by employing several signaling pathways that guarantee all regions of the genome are duplicated exactly once prior to each cell division. Inaccurate genome duplication, including partial or complete over- and under-genome duplication and other forms of replication stress, can trigger deleterious consequences such as chromosomal aberrations, stem cell failure, and the development of malignancy^1–3^.

As cells emerge from mitosis, chromosomes begin a serial recruitment process resulting in the formation of a protein complex (known as the pre-replication complex, or the replication licensing complex) that includes helicases and accessory proteins essential for genome duplication^4, 5^. The loaded helicase is inactive during the first growth (G1) phase of the cell cycle prior to the onset of DNA synthesis. During the synthesis (S) phase, cyclin-dependent kinases activate the helicase and facilitate the recruitment of additional components^4^ that allow the helicase to unwind DNA, possibly by forming a phase-separated molecular machine^6^. On each chromosome, replication starts at distinct regions termed replication origins, which initiate replication sequentially during S phase. DNA synthesis initiates at replication origins once the helicases incorporated into pre-replication complexes are activated, forming active replication forks ^4, 7, 8^. As the helicases are activated, the pre-replication complex proteins responsible for recruiting helicases to chromatin are degraded, preventing the further recruitment of helicases to the genomic regions already undergoing duplication^9–12^.

In metazoans, pre-replication complexes are bound to chromatin in excess, but chromosome duplication initiates only at a fraction of the many potential origins. The selection of active origins among the excess of pre-replication complex binding sites is often tissue-specific^8^ and preferentially associates with specific chromatin modifications ^8, 13^. The excess potential origins (“dormant origins”), which are bound by pre-replication complexes but do not initiate replication during normal mitotic growth, can be activated under distinct conditions that slow or stall replication fork elongation (replication stress^1^). This observation suggests that the recruitment of pre-replication complexes to flexible or consistently dormant origins might serve as a redundancy to ensure the prevention of incomplete genome duplication. The flexible choice of replication origins could also preserve genome integrity by coordinating replication with transcription and chromatin assembly on the shared chromatin template^8^.

CDT1, a key component of the pre-replication complex, interacts with a six-protein complex (ORC – Origin Recognition Complex) and an additional adaptor protein, CDC6, to recruit the replicative helicase (MCM) to chromatin^4, 14–16^. As cells progress through the cell cycle, multiple mechanisms are employed to facilitate CDT1 inactivation and degradation; interactions with the cell cycle regulator geminin^10, 12^ inactivate CDT1, and several ubiquitin ligase complexes, including CRL1/SCF coupled with SKP2^10, 17^ and FBXO31^18^ and CRL4 coupled with CDT2^9, 19, 20^, target CDT1 for degradation. The modulation of CDT1 levels to prevent the over-recruitment of pre-replication complexes is critical for genomic stability, as faulty degradation of either CDC6 or CDT1 can induce tumorigenic transformation via oncogene-triggered re-replication^3, 21, 22^, abnormal stem cell proliferation^23^, and sensitivity to DNA damaging agents^24^. Degradation of CDT1 is coupled to cell cycle progression in proliferating cells by an interaction between the CRL4 ligase, which degrades CDT1, and the DNA polymerase accessory ring Proliferating Cell Nuclear Antigen (PCNA)^15^.

Our previous studies have shown that RepID, a replicator-specific binding protein essential for site-specific initiation of DNA replication in mammalian cells^25^, recruits CRL4 to chromatin in a PCNA-independent manner prior to the onset of S phase and promotes CDT1 degradation. RepID deficiency is associated with delayed CDT1 degradation, resulting in limited genome re-replication^26^. Partial genome re-replication can also be caused by inhibition of DOT1L, a methyltransferase which catalyzes the demethylation of histone H3 on lysine 79 (H3K79Me2), thereby removing a histone modification typically associated with a group of replication origins^27^. Because massive re-replication can drive cell death specifically in checkpoint-compromised cancer cells, both CDT1 stabilization by inactivation of ubiquitin-mediated degradation and inhibition of DOT1L are currently being explored as novel anti-cancer therapeutic strategies ^28–33^.

Given that genome re-replication is a common avenue to genomic instability and considering its potential as a strategy for chemotherapy, it is important to understand in detail its mechanics and downstream consequences. Here, we report that re-replication exhibits aberrant replication fork dynamics and occurs more slowly than the routine genome duplication that takes place during normal growth. The consequences of the persistent presence of pre-replication complexes on chromatin include massive DNA damage and the induction of senescence. Unlike other instances of replication origin over-activation, such as when replication slows down as a result of nucleotide depletion or in cells exposed to DNA damaging conditions, the re-replication process preferentially utilizes a subset of the replication origin pool typically used during normal growth.

## Results

### CDT1 stabilization leads to massive genome re-replication and altered replication fork dynamics

To determine replication dynamics in cells undergoing replication during normal mitotic growth and in cells undergoing re-replication, we first sought to establish an experimental system in which the majority of cells undergo re-replication. To that end, we have explored several avenues for triggering re-replication. Consistent with our previous observations, partial genome re-replication was triggered by inhibiting SKP2, a key component of CRL1, in a RepID-deficient cell background^26^ (Supplementary Fig. 1a top).

RepID is a component of the CRL4 complex and both CRL1 and CRL4 complexes are crucial to prevent DNA re-replication. In both RepID-proficient and RepID-deficient backgrounds, exposure to the NEDDyation inhibitor MLN4924 (pevonedistat) — a drug currently undergoing clinical trials and known to inhibit both CRL1 and CRL4 — efficiently triggered massive re-replication in a large fraction of the cell population. For example, exposure of HCT116 cells to a very low dose of 31.25nM MLN4924 for 48h triggered partial re-replication while higher doses of 125 and 250nM MLN4924 resulted in re-replication in nearly all the cells (Fig. 1a). In cells exposed to moderate levels of MLN4924, re-replication was detected as early as 8h after addition of the drug (for example, 7.65% over-replicating >4N cells in 250nM MLN4924 treated HCT116 cells vs 1.87% in control cells) while nearly 80% of all cells were undergoing re-replication after 48h (Fig. 1b). Similar results were observed in RepID wt and RepID depleted U2OS cells (Supplementary Fig. 1a, bottom and 1b). Another NEDDylation inhibitor, TAS4464, also triggered re-replication at a lower concentration for 24h (Supplementary Fig. 1c). Because exposure to MLN4924, a drug currently in clinical trials, caused the highest fraction of cells to undergo genomic re-replication, this drug was selected as the pharmacological study.

**Fig. 1.**
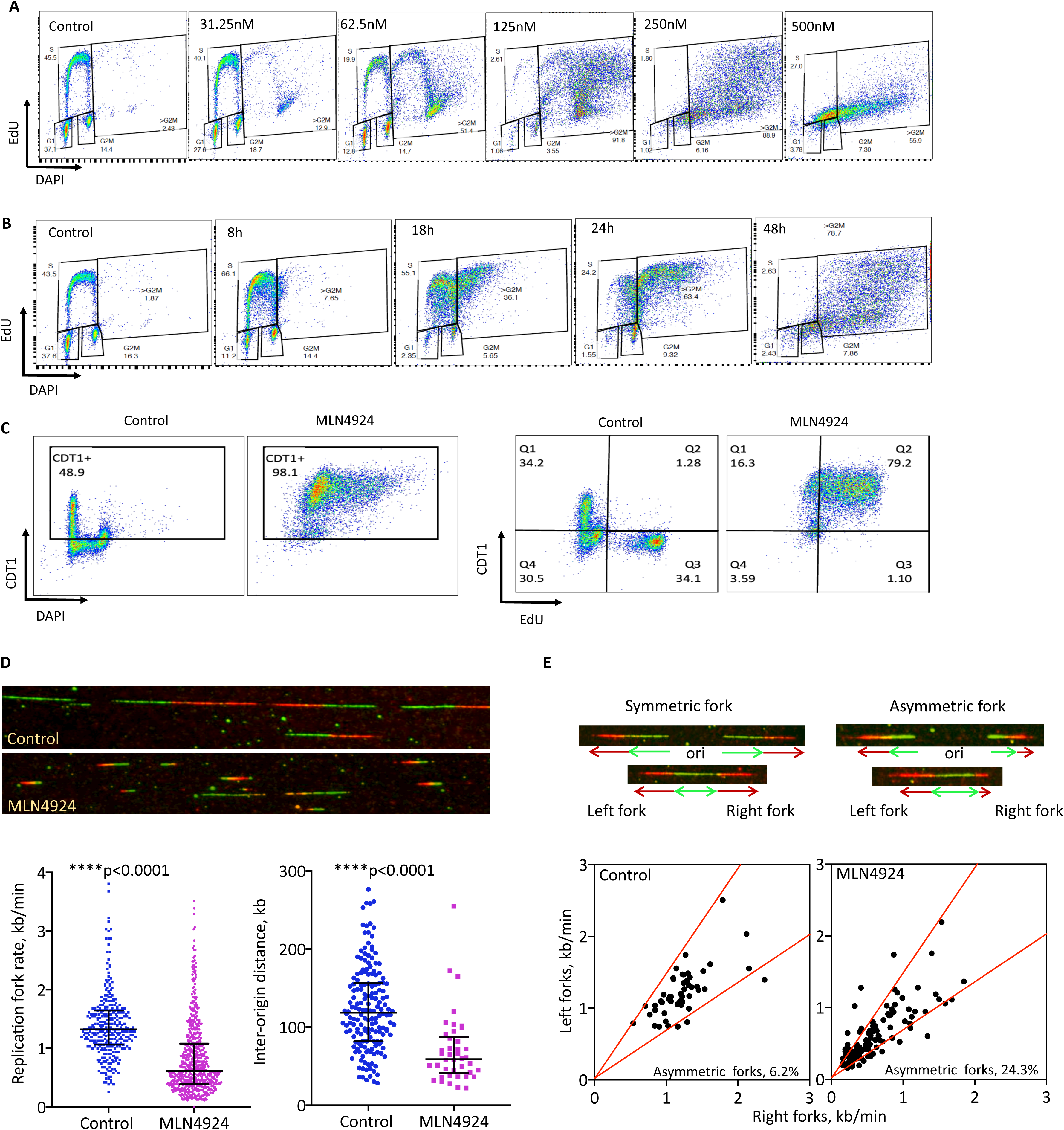
Stabilization of pre-RC components induces massive re-replication. **a,** Cell cycle profiles of HCT116 cells treated with the indicated doses of MLN4924 for 48h, or **b**, with 250 mM MLN4924 for different time periods. **c**, HCT116 cells were treated with 250nM MLN4924 for 24h. Changes in CDT1 levels corresponding to cell cycle are indicated by DAPI-DNA (left) and by the fraction of CDT1 and EdU double positive cells (right). **d**, Replication profile changes in HCT116 cells treated with 250nM MLN4924 for 30h measured by DNA combing. Cells were labeled with the thymidine analog IdU for 20 minutes (green), then CldU (red) for 20 minutes before collecting. Representative images of DNA combing for both the control and MLN4924-treated samples (top) are shown. The replication fork progression rates (bottom left) and replication inter-origin distances (bottom right) are indicated. Mann-Whitney test was used for the statistical significance. **e**, MLN4924 induced asymmetric replication forks in HCT116 cells. Cells were treated as in Fig. 1d. The lengths of left forks and right forks emanating from the same origins were compared, and if the difference was more than 30%, then they are classified as asymmetric forks. The percentages of asymmetric forks from each group are shown.

NEDDylation inhibitors such as MLN4924 inhibit CRL4 and CRL1 ubiquitin ligase complexes that target the pre-replication complex component CDT1 for degradation. In accordance with this mechanism, as shown in Fig. 1c, CDT1 levels in cells undergoing normal mitotic growth were high during the G1 phase of the cell cycle (left panel), but were very low in S phase (EdU-positive cells, right panel). CDT1 levels increased, but still remained quite low, during G2M phase, consistent with a role of CDT1 in mitotic kinetochores^15, 34^. In contrast, CDT1 levels were very high in almost all cells treated for 24h with MLN4924 (left). Most MLN4924-treated cells were EdU-positive (right), suggesting that CDT1 accumulation in MLN4924-treated cells that would promote re-licensing and re-replication. Immunoblotting confirmed increased CDT1 level starting as early as 4h of MLN4924 treatment in HCT116 cells (Supplementary Fig. 1d). Levels of both the CRL4 targets CDT1 and p21 were also higher in TAS4464 treated cells (Supplementary Fig. 1e), suggesting that both neddylation inhibitors, MLN4924 and TAS4464, induced re-replication by stabilizing the pre-RC component CDT1.

To characterize replication fork dynamics in re-replicating cells, we sequentially labeled MLN4924-treated and control HCT116 cells with IdU for 20 minutes, followed by CldU for 20 minutes. Cells were harvested, and DNA was isolated and subjected to molecular combing to detect IdU (green) and CldU (red)^35^. DNA combing (representative images shown in Fig. 1d, top panel) showed that MLN4924-treated cells exhibited significantly slower replication and shorter inter-origin distances (Fig. 1d, bottom panel), suggesting a significant increase in the frequency of replication initiation. We also observed an increased frequency of asymmetric forks in MLN4924-treated cells (Fig. 1e), suggesting that replication forks stalling at a high frequency in re-replicating MLN4924-treated cells.

Since NEDDylation inhibitors act indiscriminately on E3 ubiquitin ligases and could therefore lead to pleiotropic effects, we next looked at replication dynamics in CDT1 over-expression (CDT1-OE) cells. Stable over-expression of CDT1 was toxic to cells and no stable CDT1-OE clones could be established. We therefore established cell lines in which CDT1-OE could be triggered by exposure to doxycycline (DOX) (Fig. 2a, left panel; CDT1 was tagged with GFP, and CDT1 levels were monitored by flow-cytometry). In such cells, exposure to doxycycline for 24h resulted in the accumulation of cells with 4N DNA content, and exposure for 48h resulted in genome re-replication in most (64.9%) cells (Fig. 2a, middle panel). In addition to the increased DNA content, we noted that CDT1-OE also increased cell size and granularity (Fig. 2a, right).

**Fig. 2.**
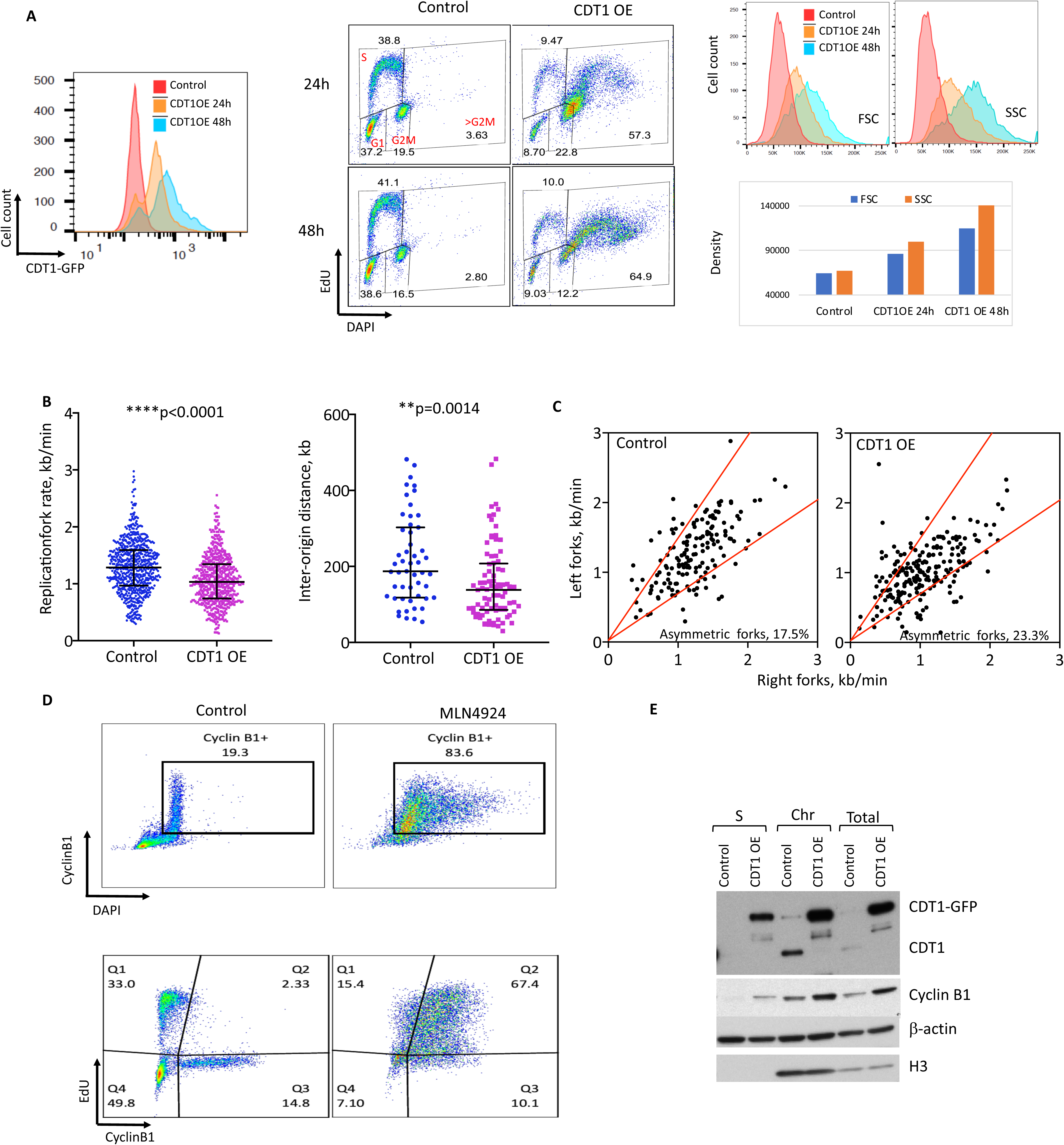
CDT1 OE induced re-replication accompanied by altered replication fork dynamics. A doxycycline inducible CDT1-GFP plasmid was stably transfected into U2OS cells (inducible CDT1-U2OS cells). **a**, Inducible CDT1-U2OS cells were cultured without (control) or with (CDT1 OE) doxycycline for 24 and 48h. CDT1-GFP expression levels (left), cell cycle progression (EdU indicating DNA synthesis, DAPI indicating DNA content, middle panel), cell size (forward scatter: FSC) and cell granularity (side scatter: SSC, right panel) were measured by flow cytometry. **b**, DNA replication profiles of inducible CDT1-U2OS cells with and without doxycycline for 48h were determined using DNA Combing. DNA replication fork progression rates (left), inter-replication origin distances (middle). Mann-Whitney test was used for the statistical significance. **c**, Asymmetric replication forks in inducible CDT1-U2OS cells treated with doxycycline for 48h. Percentage of asymmetric forks from each group are shown. **d**, HCT116 cells were treated with or without 250nM of MLN4924 for 24h and cyclin B1 levels were analyzed by flow cytometry. **e,** Immunoblots measuring changes of CDT1 and cyclinB1 levels in inducible CDT1-U2OS cells treated with doxycycline for 48h. S: cytoplasmic fraction; Chr: chromatin enrichment fraction after pre-extraction with 0.25% NP40 in low salt solution; Total: whole cells lysate.

Similar to the kinetics observed after exposure to NEDDylation inhibitors, increases in CDT1-GFP levels and re-replication were both detected as early as early as 8h after continuous exposure to doxycycline (Supplementary Fig. 2a). Notably, cells that lost CDT1-GFP expression did not undergo re-replication in the presence of doxycycline (Fig. 2a, Supplementary Fig. 2b). Single-fiber analyses demonstrated that CDT1-OE, similar to MLN4924 treatment, triggered slow replication, short inter-origin distances, and asymmetric replication forks (Fig. 2b, c).

The increased cell size suggested that cells with increased CDT1 levels continued to grow while not undergoing mitosis. To test this hypothesis, we measured the levels of cyclin B1, which starts accumulating during mid-G2 phase in normal mitotic cells and reaches its peak at the early/mid-M phase. Cyclin B1 is targeted for degradation during mitosis by the anaphase-promoting complex (APC)^36^. A flow-cytometry analysis of cyclin B1 levels in HCT116 cells showed that in untreated cells, cyclin B1 levels were very high in the G2/M phase, and all the cyclin B1 -positive cells were EdU-negative (Fig. 2d, control). However, in the MLN4924-treated HCT116 cells, the proportion of cyclin B1-positive cells significantly increased (83.6% in MLN4924-treated vs. 19.3% in control); surprisingly, most cyclin B1-positive cells were shown to be replicating (67.4% of MLN4924-treated cells, EdU-positive vs. 2% in control, Fig. 2d). The elevated level of cyclin B1 was confirmed by immunoblotting (Fig. 2e). To evaluate the frequency of mitotic cells, we used phospho-H3(Ser10), a mitotic phase cell marker^37^, in cells treated with 250nM MLN4924 for 18h or 24h. As shown in Supplementary Fig. 2 c and d, the fraction of mitotic cells was very low as early as 18h after the onset of MLN4924 treatment, suggesting that MLN4924-treated cells skipped mitosis, re-licensed replication origins, and over-replicated genomic DNA regions.

### Replication stress triggered by re-replication leads to senescence

As shown above, increased CDT1 levels and DNA re-replication are associated with decreased replication fork progression together with increased frequency of asymmetric replication. These observations often indicate replication stalling and DNA damage. To assess the consequences of re-replication in CDT1-OE cells, we measured the levels of gamma-H2AX (γH2AX) and phospho-RPA (p-RPA), which are markers for DNA double-strand breakage and replication stress, respectively. Indeed, both U2OS and HCT116 cells treated with MLN4924 show significant increased γH2AX and p-RPA levels (Fig. 3a, b). Similar results were obtained in cells harboring the inducible CDT1 construct treated with DOX for 48h or exposed MLN4924 (Supplementary Fig. 3a, b). We also noted that both MLN4924 treated cells and CDT1 over-expressing cells phosphorylated Chk1 kinase (a marker for DNA damage checkpoint activation), and γH2AX were elevated in whole cell protein extracts obtained from U2OS cells overexpressing CDT1 or treated with MLN4924 for 1 day and 2days (Fig. 3b). DNA damage levels increased quickly after CDT1 OE induction with p-RPA and γH2AX showing abnormal levels 6 h after CDT1 induction (CDT1 OE appearing 2 h after DOX treatment compared to 8 h for DNA damage) (Fig. 3c). Long-term treatments for up to 6 days showed transient p-RPA response with levels peaking 2 days after the onset of CDT1 induction and decreasing the following days (Supplementary Fig. 3b top). This observation is paralleled to EdU incorporation kinetics and could point to transient replication stress peaking at day 2 in re-replicating cells (Supplementary Fig. 3b bottom). In contrast, levels of γH2AX continuously increased throughout the 6 day testing period, indicating the constant presence of unrepaired DNA damage in the cells (Supplementary Fig. 3b).

**Fig 3.**
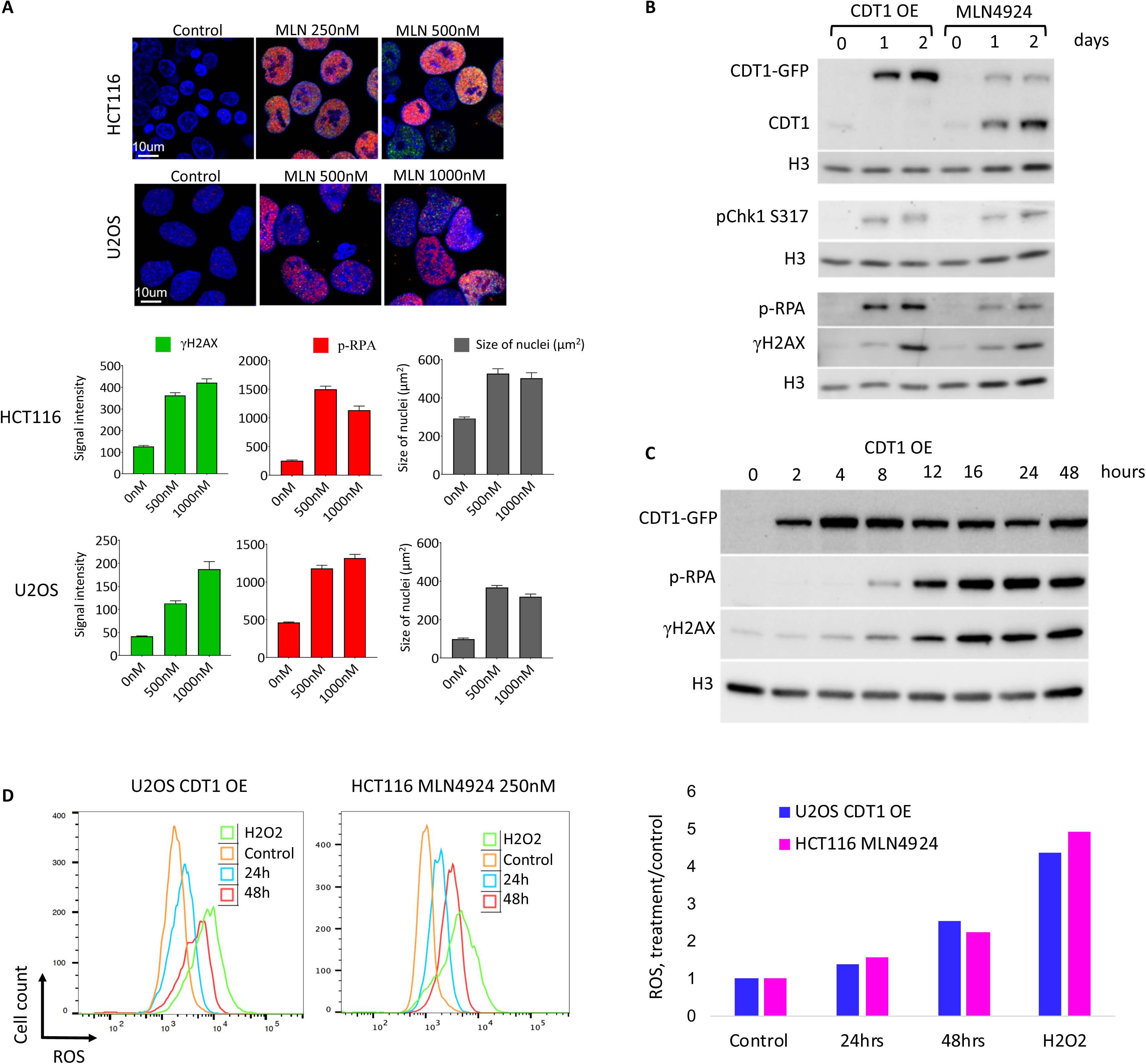
DNA re-replication induces DNA damage and activates the DNA damage checkpoint. **a,** Representatives images of HCT116 and U2OS cells were treated with the indicated doses of MLN4924 for 48h (*top panel)*,. Cells were then stained with p-RPA (red) or γH2AX (green). DNA was counterstained with DAPI. *Bottom panel*, Fluorescence intensities (arbitrary units) for p-RPA or γH2AX and nuclei size (µm^2^) are shown. **b,** Inducible CDT1-U2OS cells were treated with doxycycline or 500nM of MLN4924 for 1 day and 2 days and changes in p-RPA, γH2AX and pChk1 S317 levels were monitored by immunoblotting. Histone H3 was used as loading control. **c,** U2OS cells with inducible CDT1 OE were treated with doxycycline for 2h to 48h and changes in levels of p-RPA, γH2AX and CDT1 were monitored by Western. Histone H3 was used as loading control. **d**, Cells were treated with 1µg/ml doxycycline (CDT1-U2OS, left) and 250nM MLN4924 (HCT116, right) for the indicated times then and incubated with the ROS Deep Red Dye probe for 1 h prior flow analysis. Flow cytometry was performed immediately after harvesting cells to measure ROS levels. As a positive control, 1mM of H2O2 was added to cells together with the ROS Deep Red Dye probe. ROS levels (left) and mean fluorescent density of ROS normalized to control (right) are shown.

These results suggested that re-replication caused replication stress and RPA phosphorylation. Retained increasing γH2AX levels were indicative of a subsequent event to replication stress and RPA phosphorylation. To assess whether RPA phosphorylation and subsequent increased γH2AX levels were caused by CDT1-OE induced replication stress, we used serum starvation for 3 days to reduce the overall number of cells present in S phase, then added doxycycline to these cells for 24 h. Flow cytometry analysis confirmed that FBS starvation reduced more than 3 times the percentage of cells in S phase (50% in FBS containing medium *vs*. 15.2% in FBS-deprived medium) (Supplementary Fig. 3C, left panel). Re-replication was also significantly reduced in serum-starved cells (69.9% *vs.* 19.9% in control cells). Although CDT1 levels increased with similar kinetics in both control medium and FBS-deprived medium, both p-RPA and γH2AX levels were dramatically reduced in FBS-deprived treated cells (Supplementary Fig. 3c, right). This result demonstrates that DNA synthesis, but not CDT1 overexpression itself, is responsible for DNA replication stress and DNA damage.

As DNA replication stress often triggers reactive oxygen species (ROS) (ref), we compared ROS levels in normal and re-replicating cells. Thus, we measure ROS levels in U2OS cells expressing inducible CDT1 construct untreated and treated with doxycycline for different time periods. As a positive control, H2O2 1mM was added to cell cultures 1 h prior to ROS detection by flow cytometry. Significant increases in ROS levels were observed in re-replicating cells with maximum levels occurring 48 h after CDT1 induction (Fig. 3d, left). Importantly, ROS were induced in cells producing CDT1-GFP but not GFP alone (Supplementary Fig. 3d). Accrued ROS production was also observed in cells treated with MLN4924 (Fig. 3d, right). These results suggested that ROS may play a role in the sustained re-replication-induced DNA damage observed several days after CDT1 induction.

One outcome occurring with sustained ROS generation and DNA damage is cellular senescence (ref). To test if re-replicating cells underwent senescence, CDT1-OE inducible cells were treated with doxycycline or with 500nM of MLN4924. After 6 days, cells were tested for the cellular senescence marker β-galactosidase. As shown in Fig. 4a, both doxycycline and MLN4924 treated cells were visible as large re-replicating cells with strong blue staining. In contrast, cells with inducible GFP-only did not undergo senescence and did not exhibit increased p-RPA and γH2AX levels (Fig. 4a, b). Increase in mitochondrial mass, another hallmark of senescence, were also observed several days after re-replication (Fig. 4c). These results suggest that replication stress induced by elevated CDT1 levels eventually triggered ROS and led to cell senescence.

**Fig 4.**
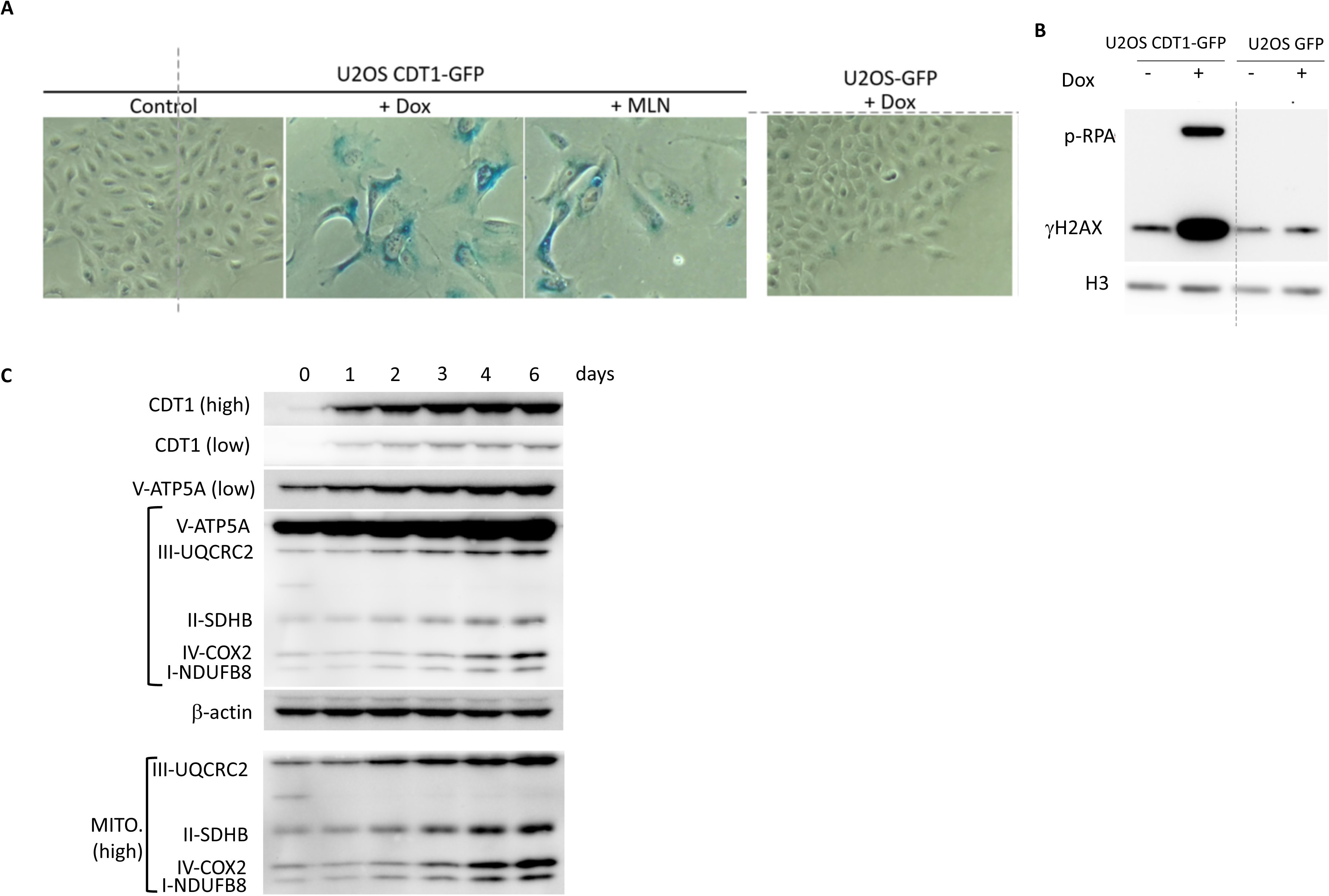
DNA re-replication induced by increased CDT1 level leads to senescence. a, Inducible CDT1-GFP-U2OS cells were treated with 1µg/ml doxycycline or 500nM of MLN4924 for 6 days, then stained with β-galactosidase to detect senescence. Inducible GFP-U2OS cells treated with 1µg/ml doxycycline for 6 days as control. b, Inducible CDT1-GFP U2OS cells and inducible GFP U2OS cells were treated with 1µg/ml doxycycline for 48h and whole cell extracts were tested for p-RPA and γH2AX level by Western. Histone H3 was used as loading control. c, Mitochondria mass in CDT1-OE cells was determined by analyzing the expression of 5 mitochondrial proteins by Western for up to 6 days of doxycycline treatment. Beta actin was used as loading control. High, high exposure; low, low exposure.

### Genomic distribution of re-replicated DNA

We next used a BrdU/CsCl gradient (a variation of the Meselson-Stahl assay^38^, outlined in Fig. 5a) to isolate re-replicated DNA in cells undergoing partial genome re-replication. HCT116 cells were labeled with BrdU for 14h, a timeframe that allowed cells to complete a single round of DNA replication in the presence of BrdU in mitotically growing cells. This timeframe was expected to enable BrdU substitution in a single DNA strand without allowing cells to complete mitosis and initiate a second round of DNA replication that would have resulted in BrdU substitution in both DNA strands. As a control, exponentially growing cells were labeled with BrdU for 48h (resulting in BrdU substitution in both DNA strands). Following the BrdU labeling, genomic DNA was fragmented and fractionated using a CsCl gradient. The fractions corresponding to completely substituted DNA (both strands containing BrdU) were sequenced. We detected DNA substituted with BrdU on both strands (heavy-heavy DNA) in cells exposed to MLN4924 (200nM and 400nM) for 14h (Fig. 5b) as well as 200nM MLN4924 for 17h and 24h (not shown). Although we did not detect very high, localized peaks, the re-replicated DNA was not distributed evenly throughout the genome and was preferentially localized to regions featuring open chromatin (Fig. 5c, d).

**Fig 5.**
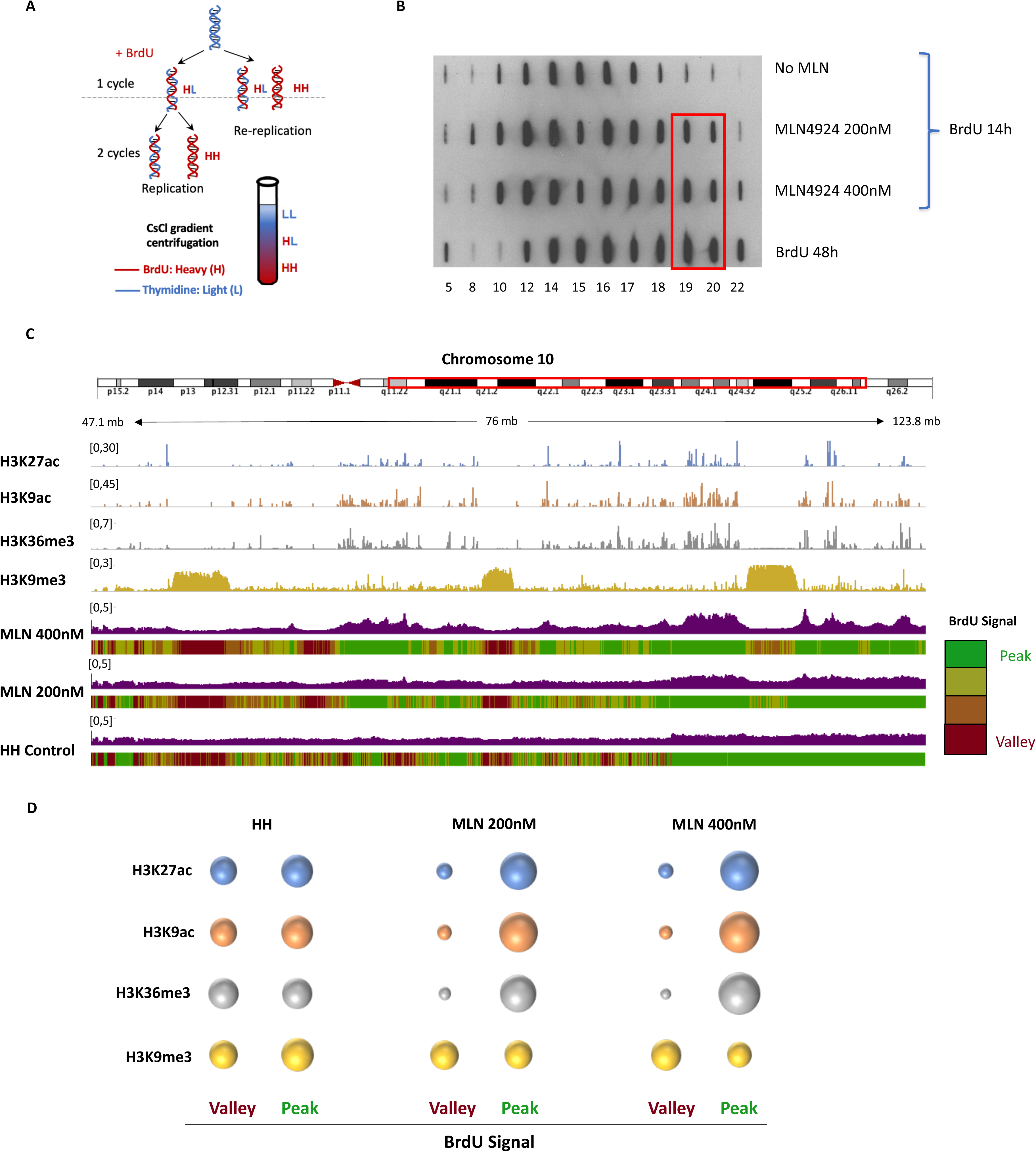
Skewed distribution of re-replicated DNA. a. Schematic representation of the method used to sequence re-replicating DNA. HCT116 cells were treated with 0, 200 or 400nM MLN4924 for 14h. BrdU was added to all samples (including the control) together with MLN4924 to trace replicating DNA. Cells incubated with BrdU for 48h, which have BrdU incorporated in both strands of DNA, were used as a positive control for re-replication. Genomic DNA was isolated and sonicated to about 0.5-1 kb, and then fractionated by CsCl gradient centrifugation. **b**, BrdU levels from equal volumes of selected fractions, from low density l (left) to high density l (right), were detected using an anti-BrdU antibody. Fractions 19 and 20 from MLN4924-treated and positive control cells (inside the red rectangle), which have BrdU incorporated into both DNA strands, were combined and sequenced. **c**, An IGV screenshot of re-replicating DNA showing the genomic location of re-replication. Tracks for H3Kk27Ac, H3K9Ac, H3K36Me3 and H3K9Me3 are from the Encode database (Bernstein Group, Broad Institute) and are used here to locate open and closed chromatin. The sequenced genome was equally divided into 4 groups according to the peak height (Green for high peak density and red for low peak density). The very wide peaks seen after MLN4924 treatment are indicative of open chromatin regions. **d**, Bubble chart showing the association of the re-replication most enriched genomic regions (top 25%, peak)) and the re-replication least enriched genomic regions (bottom 25%, valley) with heterochromatin and euchromatin, respectively. Raw data for the bubble plot and additional marks are listed in supplementary excel file 1.

The slow replication fork progression rates and the shorter inter-origin distances observed in re-replicating cells suggested that re-replicating cells might utilize a subset of origins not activated during normal growth. To test this hypothesis, we first isolated re-replicated DNA in MLN4924 treated HCT116 cells by BrdU-CsCl gradient (MLN4924-HH, both DNA strand had BrdU incorporated). DNA with only one strand with BrdU incorporation from control cells (Control-HL) and MLN4924 treated cells (MLN4924-HL) were also collected to map normal replication origins. Normal and re-replicating origins were mapped by Nascent Strand-Sequencing (NS-seq) as previously described ^39, 40^ (Fig. 6a). Combining BrdU density gradients followed by NS-seq, we sequenced only the re-replicated DNA strands that initiated replication within 2-3 minutes of DNA isolation. As shown in the representative IGV screenshots in Fig. 6b, the distribution of NS-Seq peaks in normal replication (Control-HL and MLN4924-HL) was very similar to the distribution of NS-peaks in re-replicated DNA (MLN4924-HH).

**Fig 6.**
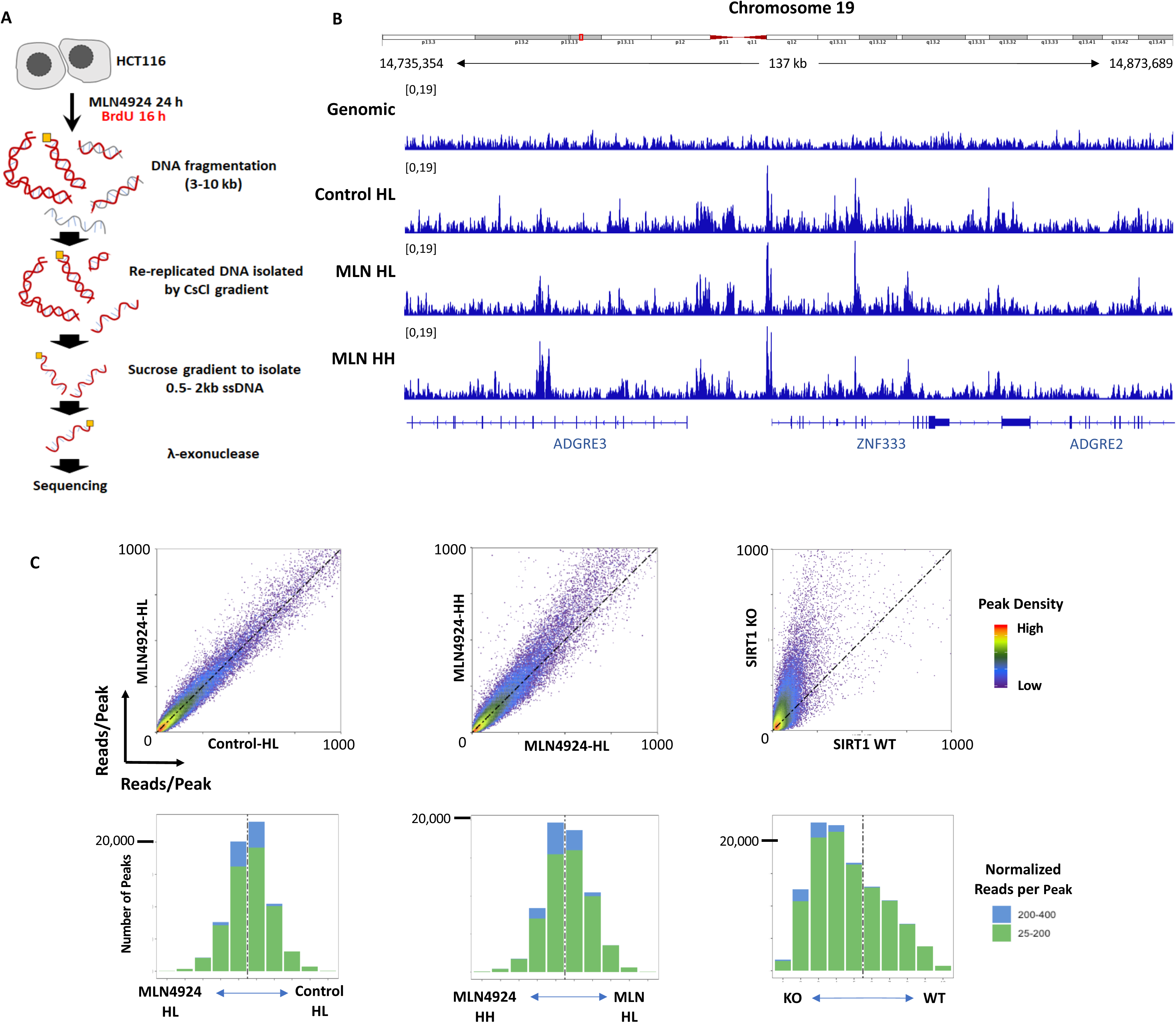
Re-replication is driven by the same set of origins utilized during normal mitotic growth. **a**, Schematic repsentation of method used to sequence re-replicating DNA. To map re-replicating origins, cells were treated with MLN4924 for 24 h, including exposure to BrdU for 16 h. Genomic DNA was fragmented and re-replicated DNA (MLN HH DNA from cells incubated with BrdU for less than an entire doubling period) was isolated by BrdU-CsCl gradient. Normal replicated DNA (Control HL DNA and MLN HL DNA, only one DNA strand labeled with BrdU) was also collected to map normal replication origins as a control. Following the CsCl gradient, newly replicated nascent DNA was isolated from both normal- and re-replicated DNA using sucrose gradient and lambda exonuclease. **b**, IGV screenshot showing representative nascent DNA outputs in genomic control, during normal mitotic replication (Control-HL and MLN-HL) and during re-replication (MLN4924-HH). **c**, Density plots (top panel) comparing replication origin usage in 2 normal mitotic replication samples (Control-HL and MLN4924-HL, left) and in one normal and one re-replication samples (MLN4924-HL and MLN4924-HH, middle). The location of each data point is proportional to the number of reads per origins. Origins that initiate replication with similar frequency during normal mitotic growth and in re-replicating cells are located on the diagonal dotted line; for the middle panel, origins above the diagonal dotted line initiate more frequently in re-replicating cells and vice versa. The right panel used as positive control for dormant origin activation showing augmented origin activation in SIRT1 depleted cells. The bar graphs below the density plots illustrate the cumulative distribution of small peaks with 25-200 reads (green) and large peaks 200-400 reads (blue).

We then compared the frequency of initiation (reads per peak) in cells undergoing normal mitotic growth with that of MLN4924-treated cells. We used peak density plots created with BAMScale (https://github.com/ncbi/BAMscale) to determine whether certain groups of replication origins are preferentially utilized during re-replication (Fig. 6c). Density plots represent peak sizes (number of reads/each peak) across the sample pairs. Similar replication initiation frequencies are reflected in similar peak sizes across the samples, with most peaks distributed along the diagonal (45 degree dotted black line in the diagram shown in Fig. 6c). Peak size distributions that are skewed towards one of the compared samples reflect a higher frequency of initiation in that sample. As shown in Fig. 6c, density plots comparing replication origin utilization in SIRT1 proficient and deficient samples clearly demonstrate that the SIRT1-deficient sample exhibited a higher level of replication origin utilization as reported^8^. In contrast, the MLN4924-treated samples did not exhibit a similar bias (Fig. 6c), confirming that dormant origins were not overwhelmingly activated in re-replicating cells.

Similar results were obtained when we used a different strategy to map re-replication origins. For this, HCT116 cells were exposed to 250nM MLN4924 for 30h or 45h, timepoints at which nearly all the replicating cells (30h) or all the replicating cells (45h) underwent genome re-replication (Supplementary Fig 4a, left panel). The cells were collected and nascent strand DNA was prepared and sequenced as described previously (NS-Seq^39, 40^; Supplementary Fig. 4a, right panel). DNA from cells not exposed to MLN4924 was used as a control. As shown in the representative IGV screenshots in Supplementary Fig. 4b, the distribution of NS-Seq peaks in the control and MLN4924-treated samples was very similar. As shown in the density plots, most re-replicating origins colocalized with the replication origins utilized during normal mitotic growth. Notably, this result differs when cells are exposed to conditions that are known to activate dormant origins; for example, in cells depleted of SIRT1^8^, where peaks that did not initiate replication in SIRT1-proficient cells were clearly observed (Fig. 6c, right). These observations suggested that re-replication was not accompanied by a large-scale activation of dormant origins in these samples. We therefore concluded that during re-replication, cells largely used the same set of origins as is used during normal mitotic growth.

### Early replicating origins are preferred during massive re-replication

DNA replication follows a stringent spatial order, forming distinct replication foci patterns in early, mid, and late S phase. As shown in Fig. 7a, exponentially growing cells exhibited very typical EdU foci patterns for early (ES), middle (MS), and late (LS) S phase. In contrast, the MLN4924-treated samples displayed homogeneous replication foci similar to the early-S-phase patterns, with almost no cells exhibiting late-S-phase foci. These observations suggested that although re-replication used the same pool of replication origins as the one from control cells, those re-replicating cells lost the spatial-order characteristics of normal replication.

**Fig. 7.**
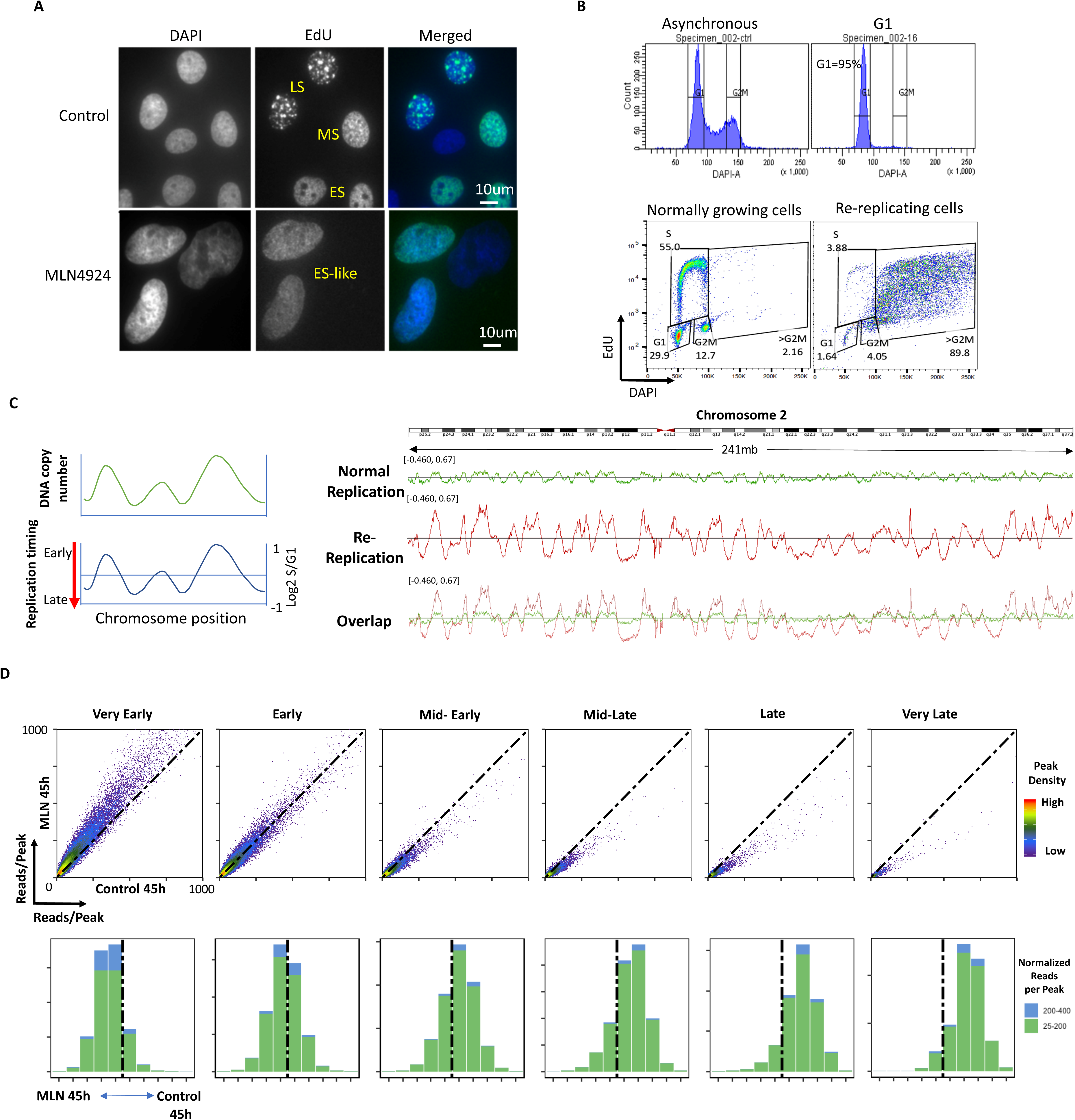
Replication origins that initiate replication early during normal mitotic growth are over-activated during re-replication. **a**, U2OS cells were treated with 500nM of MLN4924 for 48h and EdU click-it to label S phase cells. Control cells showed typical S phase cell foci patterns (ES: early S phase; MS: mid-S phase; LS: late S phase). MLN4924 treated cells lost this typical S phase replication foci patterns, showing ES-like foci patterns. **b**, Total genomic DNA of HCT116 cells was purified from G1 cells (isolated by elutriation), from asynchronous normal cell cycle cells and asynchronous re-replicating cells induced by MLN4924 treatment for 36h (when almost all the replicating cells were in a re-replicating cycle) and sequenced at >30X coverage. **c**, Since at any given time in asynchronous cells, early-replicating genomic regions are present in more copies than late-replicating genomic regions, the ratio of copy number in asynchronous cells to that in G1 cells can be used to determine the relative replication timing of any genomic region (left). IGV screenshot of replication timing of chromosome 2 for normal replication and re-replication (same y-axis scale) to show the increased copy number at the early replication genomic regions and decreased copy number at the late replication genomic regions in re-replicating cells (right). **d**, Replication origins from normal replication (control) and re-replication (MLN4924 for 45h) were classified into six groups ranging from very early to very late. Peak sizes (measured as read per peak, meaning the bigger the peak, the more frequent the initiation of the origins) of all origins in each timing group were compared between normal replication and re-replication (top panel) in dot plots. Dots (origins) on the diagonal dotted line, indicate there was the same initiation efficiency between normal and re-replication. Origins above the diagonal dotted line represent those that initiate more frequently in re-replication than in normal replication, and vice versa. *Bottom panel*, Origins in each dot plot were further divided into 10 fractions ranging from the very strong origins in re-replication (MLN 45h, left side of each bar graph) to the very strong origins in normal replication (Control 45h, right side of each bar graph) and the number of peaks (origins) with more than 400 reads (blue) and 200-400 reads (green) in each fraction are show in the bottom panel to show the shift from increased origin initiation in early S-phase to decreased origin initiation in late S phase in the re-replication sample (MLN 45hcompared to normal replication sample (control 45h).

The observation that re-replicating regions are located preferentially at open chromatin (Fig. 5c, d) and exhibit a loss of typical replication foci patterns (Fig. 7a) are consistent with the hypothesis that the early-replicating portions of the genome are over-represented in re-replicating DNA. We then determined the replication order (replication timing) of each genomic portion during normal mitotic growth and in re-replicating cells using a variation of the Timex method^41, 42^, method that is based on the assumption that in an asynchronously growing cells, early-replicating regions will be present in higher copy numbers than late-replicating regions. We determined whole-genome copy number in asynchronous, exponentially growing cells in which more than 50% of the cells were in S phase, as well as in cells in the G1 phase of the cell cycle that were isolated by centrifugal elutriation (Fig. 7b). We also measured whole-genome copy number in asynchronous re-replicating cells that were treated with 250nM MLN4924 for 36h (89.8% re-replicating cells, Fig. 7b). All samples were sequenced for more than 30X depth. The log2 value of the copy number in asynchronous replicating and re-replicating cells, normalized to cells in G1, was used to determine the replication timing (Fig. 7c left). As shown in Fig. 7c right, the copy number variations for the entire chromosome 2 are shown side-by-side with the same y-axis scale. The order of DNA replication (the location of copy number peaks and troughs) was similar in cells undergoing normal mitotic growth and in re-replicating cells. However, earlier-replicating genomic regions were over-represented in re-replicating cells whereas late-replicating genomic region were under-represented. This is consistent with the observation that re-replication preferred open chromatin regions and showed only early-S-phase-like re-replication foci patterns.

To test whether the over-represented, earlier-replicating genomic regions in re-replicating cells reflect a population of cells that had stalled replication after completing the duplication of the early-replicating portion of the genome, we directly measured the frequency of replication initiation at origins in control and MLN4924-treated samples. All origins were stratified into six groups according to replication order, ranging from very early to very late. Then, we used BAMScale to create density plots measuring the number of reads per replication origin peak during normal mitotic growth and during re-replication. As shown in Fig. 7d (top panels), very early origin peaks are mapped above the diagonal, suggesting that early replication origins initiate replication more frequently in re-replicating cells than in cells undergoing normal mitotic growth. This over-initiation was not evident in later-replicating genomic regions; in contrast, late-replicating origins initiate more frequently in cells undergoing normal mitotic growth than in re-replicating cells (Fig. 7d). Similar results were also obtained when we compared re-replication origin data obtained by the second strategy (Supplementary Fig. 5a) to replication origins mapped during normal mitotic growth. However, we did not see any origin initiation timing shifts when we compared 2 sets of normal replication samples (Supplementary Fig. 5a). Altogether, these results show that re-replication occurs throughout the genome with a prevalence towards open chromatin and early replicating regions.

## Discussion

The results reported here characterize replication re-initiation profiles in cells containing persistent pre-replication complexes (pre-RCs) on chromatin during the S phase of the cell cycle. Although pre-RCs are normally removed from chromatin following DNA synthesis, intact pre-RCs can remain on chromatin under several circumstances, including upon the over-expression of proteins responsible for preventing the degradation of key pre-RC components, such as CDT1. Experimentally, the persistence of intact pre-RCs on chromatin can be achieved by pharmacological inhibition of CRL1 and CRL4 NEDDylation or by depletion of cellular components that recruit CRL4 to chromatin (RepID). We observed that unlike normal mitotic growth, during which only a fraction of replication origins activated on each chromosome and the resulting inter-origin spacing is 100-150 kb, re-replicating cells show a higher frequency of initiation from origins positioned in closer proximity to one another. We also observed that DNA synthesis rates during re-replication are slower than DNA synthesis rates measured during normal mitotic growth, reflecting the frequent stalling of replication forks that can lead to DNA damage, ROS production and eventually trigger senescence. Surprisingly, although re-replicating cells activate replication origins at a higher frequency than cells undergoing normal mitotic growth, these additional origins are derived from the same pool of potential replication initiation sites. Another surprising finding is that the population of re-replication origins is markedly enriched for origin sequences that are normally activated during the early stages of genome duplication. A summary of our findings is illustrated in Fig. 8.

**Fig. 8:**
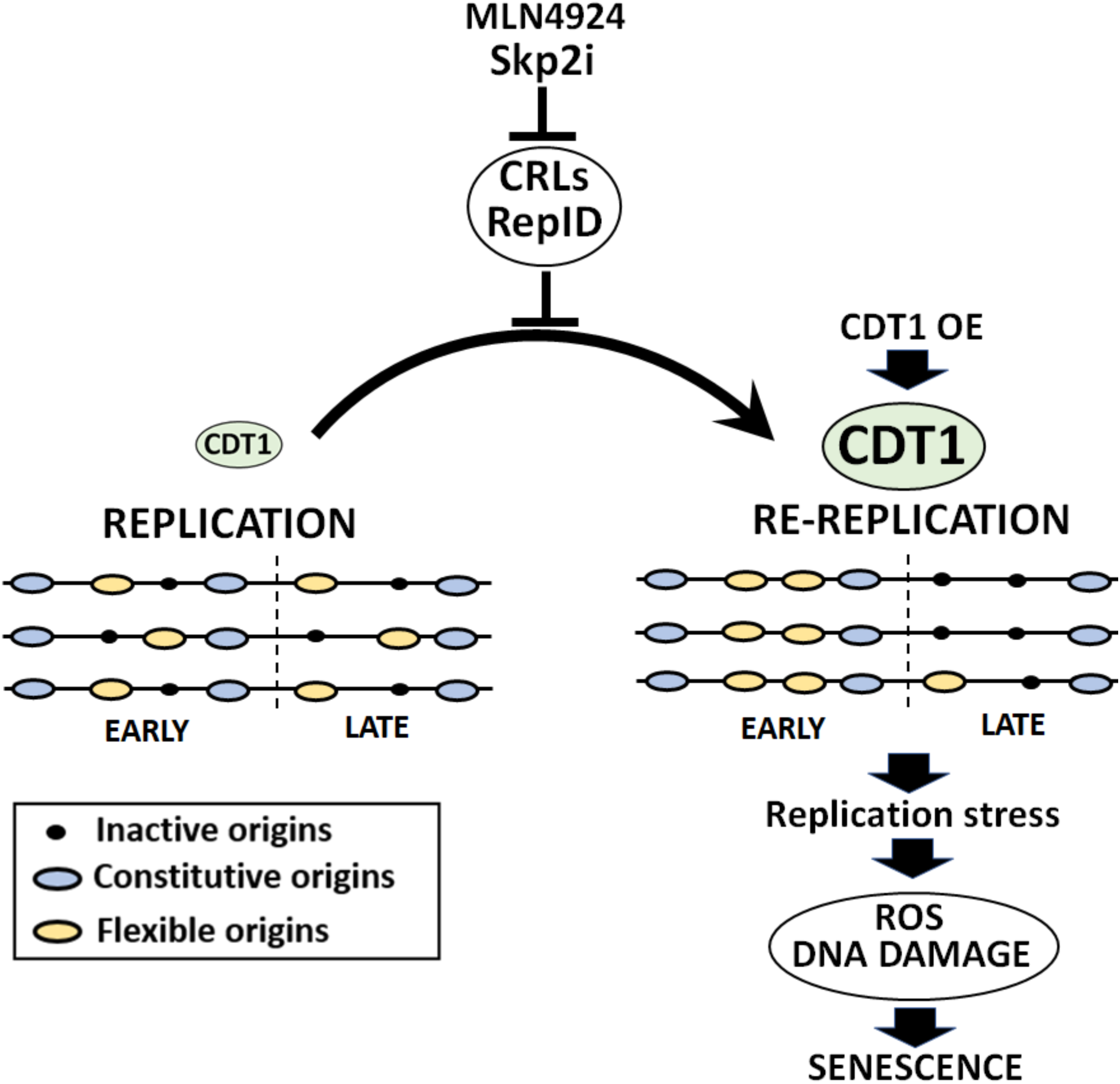
A model of replication origin usage during re-replication. Re-replication can be induced by pharmacological inhibition of the CRLs ubiquitin ligases complexes (MLN4924, Skp2 inhibitor) or by depletion of cellular components recruiting CRLs to chromatin (RepID). Both strategies lead to CDT1 stabilization and abnormal high cellular levels of CDT1. Re-replication can also be induced by induced CDT1 overexpression. Re-replication origins are markedly enriched for origin sequences that are normally activated during the early stages of genome duplication. Re-replication leads to higher frequency of origin initiation, especially flexible origins, resulting in shorter inter-origin distances. Re-replication forks are slower than those from normal mitotic growth, reflecting frequent stalling of replication forks that can lead to DNA damage, ROS production and eventually trigger senescence.

Re-replication is observed in certain developmental systems as a mechanism for controlling the development of specialized cellular functions^12, 43^. In normal somatic cells, re-replication is often fatal, activating a checkpoint control that initiate cell death to avoid carcinogenesis^44–46^. Cancer cells often trigger limited re-replication by prevention of CDT1 degradation, which is primarily catalyzed by CDT2-associated CRL4 along with an alternative degradation pathway that includes CRL1 and SKP2^9, 17, 19, 20, 31^. The interaction of CDT1 with geminin also serves to prevent re-replication during normal mitotic growth^12, 15, 17, 45, 47^. Interfering with the degradation and inactivation of CDT1, either by geminin depletion^47^ or by inhibiting the activity of cullin-anchored ubiquitin ligases^28, 31, 33, 46^, can be exploited to trigger re-initiation and subsequent selective killing of cancer cells. We observed that cancer cells that undergo partial genome duplication due to the persistence of pre-RCs on chromatin exhibit similar DNA synthesis characteristics regardless of the mechanisms that prevent pre-RC dissociation. Integrating single-fiber and genomic sequencing analyses, we found that although the frequency of initiation is higher in re-replicating cells than in cells undergoing normal mitotic growth, the initiation sites activated in both conditions are similar. These observations strongly support the hypothesis that replication origins utilized during re-replication are derived from the same set of pre-licensed origins utilized during mitotic growth.

Cancer cells can over-activate dormant replication origins when exposed to agents that slow or halt DNA replication. Because cells that undergo partial genome re-replication also exhibit slow DNA synthesis rates and undergo DNA damage, our experiments cannot unequivocally exclude the possibility that the slow replication is the cause, and not the consequence, of the activation of additional origins. We also cannot exclude the possible limited activation (below our detection threshold) of a population of origins that is dormant during normal mitotic growth. However, the population of replication origins activated in cells undergoing re-replication markedly differs from the dormant origins activated in cells undergoing over-replication due to other triggers, such as exposure to DNA damaging agents, low nucleotide pools, or depletion of SIRT1^8, 48–52^. The dormant origins activated when replication slows or stalls often include a group of DNA sequences that do not initiate replication during normal mitotic growth^1, 8, 48^, whereas the origins associated with re-replication due to the persistence of pre-RCs on chromatin are selected from the same pool of origin sequences as those activated during normal DNA synthesis.

During normal mitotic growth, the genome is duplicated in a consistent, tissue-specific order (replication timing). This order is preserved during genome re-replication but is accompanied by the preferential re-replication of regions that tend to replicate early during the normal mitotic cycle. These early-replicating origins are associated with euchromatin and therefore might represent a population of more readily accessible licensed origins. Our data show that late-replicating regions are under-represented in re-replicating DNA, suggesting that cells do not wait to complete the first round of DNA replication prior to the activation of the second set of initiation events; rather, before completing replication of the entire genome, cells initiate replication at regions that had previously undergone replication. This pattern differs from the replication pattern observed for other known instances of genome re-replication during embryonic development^43^, when cells undergo numerous iterations of entire genome duplication, with S phases and M phases alternating to re-license origins anew after the completion of each replication cycle. Because re-replicated DNA can be more susceptible to damage and breakage, the preferential re-replication of early-replicated DNA can be linked to the observed clustering of DNA breaks in euchromatic regions^53, 54^.

The existence of mechanisms that that normally select a small group of active replication initiation sites from a pool of potential replication origins can facilitate genomic stability, allowing for the complete duplication of the genome when replication stalls. Under such circumstances, excess origin activation prevents under-replication, which can lead to cell cycle perturbations, chromosomal translocations, and DNA breakage in regions with low origin density^1, 8, 50, 55, 56^. However, excess initiation of DNA replication can have deleterious consequences, including oncogenic transformation of normal cells and increased genomic instability in cancer ^28, 44, 45, 47^, consistent with our observations suggesting that massive over-replication can lead to senescence. Those consequences can be exploited therapeutically to induce selective killing of cancer cells^28, 31–33, 47^. The results reported here imply that circumventing the strong inhibitory interactions that normally prevent excess DNA synthesis can occur via at least two pathways, each activating a distinct set of replication origins. Understanding the interactions underlying both pathways could clarify the mechanisms that monitor and regulate the progression of genome duplication and lead to improved selectivity in targeting cancer.

## Methods

### Cell culture, drugs, and establishment of a CDT1 stable cell line

Human HCT1116 cells and U2OS cells were grown at 37°C in a 5% CO2 atmosphere in Dulbecco’s modified Eagle medium (ThemoFisher, 10564011), supplemented with 10% fetal calf serum. MLN4924 and SKP2 inhibitors were purchased from Cayman Chemicals (15217-1) and Millipore (506305), respectively. TAS4464 was a gift from Taiho, Inc. CDT1 cDNA was cloned by reverse transcription from HCT116 cells and verified using sanger sequencing. CDT1 cDNA was further cloned into the Tet-On 3G inducible expression system (TaKaRa, 631168) with a flag tag at the N terminal and an EGFP tag at the C terminal by the In-Fusion HD Cloning system (TaKaRa, 638909). After transfection of the U2OS cells with the pCMV-Tet3G Vector, stable clones were further transfected with the pTRE3G Vector containing flag-CDT1-GFP. Stable clones were tested for GFP positivity (CDT1-GFP) and re-replication by flow cytometry and presence of fusion protein by western blotting with both anti-CDT1 and GFP antibodies.

### Flow cytometry to monitor cell cycle, DNA re-replication and protein expression

Cells were pulse-labelled with 10 μM EdU for 30-45 min before harvest. EdU staining was performed using the Click-iT EdU kit (ThermoFisher, C10634 or C10633) according to the manufacturer’s protocol. If necessary, immunodetections with anti-cyclin B1 (Cell Signaling, 12231) and anti-CDT1 (Cell Signaling, 8064) were performed at 4 °C overnight prior to flow analysis to visualize changes in protein levels during the cell cycle. DAPI was used for DNA staining. A BD LSR Fortessa cell analyzer with FACSDiva software and/or FlowJo10.6 were used for cell cycle analysis.

### Western blotting

Most Western blots in this study were done with whole-cell lysates by adding 200μl 1 × SDS loading buffer per 1 million cell pellets. Samples were heated at 100°C for 5 minutes, centrifuged, and the supernatant was used for Western blot. In some cases, proteins were fractionated for cytosol and nuclear fractions prior to electrophoresis and immunoblotting. Cytosolic and nuclear protein fractions were collected as follows: cells were incubated on ice for 10 min in a cytosol extraction buffer (10μM HEPES, pH7.9; 10μM KCl; 1μM EGTA; 0.25% NP40; 1X protease inhibitor cocktail and phosphatase inhibitor cocktail) and, then centrifuged at 2700×*g* for 5 min at 4 °C. The supernatant was used as the “cytosol/soluble” fraction. The pellet was resuspended with 1× SDS loading buffer, heated at 100°C for 5 minutes and centrifuged, and the supernatant was used as the “chromatin-enriched” fraction for western blot analysis. The antibodies used were Phospho-Chk1 (Ser317) (Cell Signaling, 2344), anti-γH2AX (Millipore, 05-636), anti-p-RPA (Bethyl labs, A300-245A), Cyclin B1 (Cell Signaling, 12231), CDT1 (Cell Signaling, 8064), total OXPHOS human antibody cocktail (Abcam ab110411), anti-phospho-Histone H3 (Ser10) (Millipore, 05-570), anti-beta-actin (Sigma, A5316), anti-α-tubulin (Sigma, T9026) and anti-histone H3 (Millipore, 07-690).

### Microscopy

Cells were incubated with or without EdU 10 μM for 30 min, fixed with 4.0% paraformaldehyde for 10-15 min at room temperature. Cells were then permeabilized with 0.5% Triton X-100 in PBS for 30 min, blocked with 5% BSA for 30minutes and followed by Click-iT EdU labeling with/without antibody staining. EdU labeling by the Click-iT EdU kit (ThermoFisher, C10634 or C10633) was performed according to the manufacturer’s protocol. Samples were incubated for 2h with primary antibodies anti-γH2AX (Millipore, 05-636) and anti-p-RPA (1:500, Bethyl labs, A300-245A) at 1:500 dilution followed by incubation for 1h (dilution 1:500) with secondary antibodies (Alexa 488 conjugated anti-mouse IgG and Alexa 568 conjugated anti-rabbit IgG (Thermo Fisher Scientific, A11029 and A21428)). DNA was counterstained with DAPI. The Zeiss LSM710 confocal microscope and the BD pathway 855 microscope were used for imaging.

### β-galactosidase staining method

Cells were treated with doxycycline for 7 days and were fix and stained with a Senescence β-Galactosidase Staining Kit (Cell Signaling, 9860) according the manufacture’s instruction and incubated at 37 °C for overnight.

### DNA replication analysis by molecular combing

Analysis of DNA replication by molecular combing was performed as previously described ^35^. Briefly, asynchronous cells were sequentially labelled with 20 μM IdU for 20 min and 50 μM CldU for 20 min and chased with 200 μM thymidine for 60–90 min. To preserve long genomic DNA fibers, harvested cells were embedded in low melting point agarose plugs. The plugs were incubated in cell lysis buffer with proteinase K at 50°C for 16 hours, washed 3 times with TE buffer, and then melted in 0.1M MES (pH 6.5) at 70°C for 20 min. Agarose was subsequently degraded by adding 2 μl of β-agarase (Biolabs). To stretch DNA fibers, DNA solutions were poured into a Teflon reservoir and DNA was combed onto salinized coverslips (Genomic Vision, cov-002-RUO)) using an in-house combing machine. Coverslips were visually examined for DNA density and fiber length by YOYO1 DNA staining (Invitrogen). Combed DNA on coverslips was then baked at 60 °C for 2 h and denatured in 0.5 N NaOH for 20 min.

Coverslips were blocked for 10 min in 5% BSA. IdU, CldU and single-strand DNA were detected using a mouse antibody directed against BrdU (IgG1, Becton Dickinson, 347580, 1:25 dilution), a rat antibody directed against BrdU (Accurate chemical, OBT0030, 1:200 dilution) and a mouse antibody directed against single-stranded DNA (ssDNA) (IgG 2a, Millipore, MAB3034, 1:100), respectively. Incubation with primary and secondary antibodies were performed at room temperature in 1% BSA in PBS for 1 h and 45 min respectively. The secondary antibodies used were goat anti-mouse cy3 (Abcam ab6946), goat anti-rat cy5 (Abcam, ab6565) and goat anti-mouse BV480 (Jackson ImmunoResearch, 115-685-166) for ssDNA. Slides were scanned as described previously^35^ or with a FiberVision Automated Scanner (Genomic Vision). Replication signals on single DNA fibers were analyzed using FiberStudio (Genomic Vision). Only replication signals from high-quality ssDNA (not those from DNA bundles nor those located at the end of a strand) were selected for analyses. Experiments were performed at least in duplicate using independent biological isolations of DNA fibers for each experimental condition. The statistical analyses were performed using Prism 8 (GraphPad software) and the non-parametric Mann– Whitney rank sum test.

### BrdU-CsCl gradient to isolate re-replicated DNA

HCT116 cells were cultured with 50μM BrdU for the indicated times. Genomic DNA was purified and sonicated to 500–2,000 bp. Sonicated genomic DNA from cells cultured with/without BrdU for 48h were used as BrdU positive and negative control, respectively. 300 µg of sonicated DNA were fractionated at 45,000 rpm in a Ti75 rotor (Beckman) for 66h using 6 ml CsCl (1 g/ml in TE). Fractions of 250 µl were collected and the refractory index was measured to confirm the formation of the CsCl gradient. Samples of equal volume from each fraction were loaded to a positively charged nylon membrane using a Slot Blot Filtration Manifold (PR648, GE Healthcare Life Sciences, PR648). The presence of BrdU on the nylon membrane was detected with an anti-BrdU antibody. DNA in which both strands had undergone BrdU incorporation was collected and sequenced using the Illumina genome analyzer II.

### Strategy 1, performing NS-seq in re-replicating cells

HCT116 cells were treated with 250nM of MLN4924 for the indicated time in order to have most or all cells in re-replicating cycle. Cells incubated without MLN4924 were used as control. Genomic DNA was purified, and nascent strand were isolated as performed as described previously^39, 40^. Briefly, DNA was denatured by boiling for 10 min, immediately cooled on ice, and fractionated on a neutral sucrose gradient. Fragments, 0.5–2 kb, (containing nascent strand DNA and broken genomic DNA) were collected and treated with λ exonuclease to remove non-RNA-primed broken genomic DNA. Remaining single stranded nascent strand DNA was converted to double-strand DNA using the BioPrime DNA Labelling System (ThermoFisher, 18094011). Double-stranded nascent DNA (1 μg) was sequenced using the Illumina genome analyzer II (Solexa). Sheared genomic DNA was also sequenced to be used for peak calling.

### Strategy 2: isolating re-replicated DNA followed by NS-seq

HCT116 cells were treated with MLN4924 for 24h. BrdU was added to cells 8h after MLN4924 treatment for a total of 16h of BrdU incorporation, which was less than one doubling time. DNA was purified from these cells and sonicated to 3 to 10 kb. Re-replicated DNA (in which both DNA strands had undergone BrdU incorporation) was isolated using a BrdU-CsCl gradient. Re-replicated DNA and some DNA that had not undergone re-replication (only one strand having incorporated BrdU) were collected for NS-seq as described below.

### Replication timing in normal and re-replicating cells

HCT116 and U2OS cells were treated with MLN4924 for the indicated times and doses. Untreated G1 cells were isolated by elutriation at 2000 rpm, at 4°C, at flow rate 15 ml/minute for HCT116 cells, 20ml/minute for U2OS cells. DNA from both G1 (>98% in G1 phase) and exponential growing (>50% cells in S phase) untreated cells as well as from re-replicating cells were extracted using the Qiagen DNeasy blood and tissue kit (cat# 69581). DNA samples were pooled and sequenced on HiSeq using Illumina TruSeq Nano DNA library preparation and paired-end sequencing. The mean coverage was at least 30X depth.

### NGS analysis

Raw FASTQ sequencing files were first trimmed with the *Trimmomatic* (version 0.36) ^57^ and *Trim Galore* (version 0.4.5)^57^ programs to remove low quality reads. Trimmed FASTQ files were then checked for quality using *FastQC* (version 0.11.5) [https://www.bioinformatics.babraham.ac.uk/projects/fastqc/]. Trimmed reads were aligned to the hg19 genome using the *bwa* aligner (version 0.7.17) ^58^. Peaks with high read coverages were identified by the narrow *MACS2* (version 2.1.1.20160309) ^59^ peak calling method. Peaks were filtered using the “peak-score” *MACS2* metric in R (version 3.5.1) by accepting regions above the inflection-point threshold of “peak-scores” from the raw output. In order to compare samples by coverage, the “BAMscale cov” method was prepared with merged nascent-strand regions and alignment files for each sample. Regions were assigned normalized coverage values based on the library size normalization method of *BAMscale*. Peak density plots comparing sample pairs were created using R, the code is available at the *BAMscale* GitHub page (https://github.com/ncbi/BAMscale/wiki). For viewing in the genome browser, the *BAMscale* “scale” method was used to develop scaled bigwig coverage tracks for each alignment file in the using the GIGGLE search engine. Post-calculation analyses included the development of an inclusion ratio:

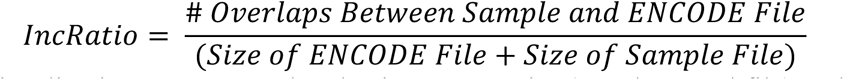

Subsequent visualizations were completed using an R-script (supplemental file) and Excel.

For BrdU-CsCl assay to detect the distribution of re-replicated DNA, samples were analyzed using the bigwig segmentation R-script available of the *BAMscale* GitHub page. Coverage files were separated into quartiles for “lower (valley)” to “upper (peak)” and visualized using IGV. The chromatin features of these re-replication peaks and their surrounding genomic regions up to 35kb were analyzed.

Replication timing data was processed by the sequencing facility using the *DRAGEN* analysis pipeline (01.003.044.02.05.01.40152). Data received from the facility was then transformed into log2 ratio coverage tracks using the “BAMscale scale” method and associated “ --operation log2” flag. Separation of replication timing regions from every early to very late was completed using the log2 ratio of re-replication vs G1 (percentile of total ratio-range: 0-10% for Very Early; 10-30% for Early; 30-50% for Mid-Early; 50-70% for mid-Late; 70-90% for late and 90-100% for Very Late). R-scripts which were utilized for this task are available in the *BAMscale* GitHub page.

## Acknowledgements

This study was supported by the Intramural Research Program of the NIH, Center for Cancer Research, National Cancer Institute. We thank the CCR core sequencing facility headed by Bao Tran and Jyoti Shetty for expert help with DNA sequencing.

---

**Supplementary Fig. 1.**
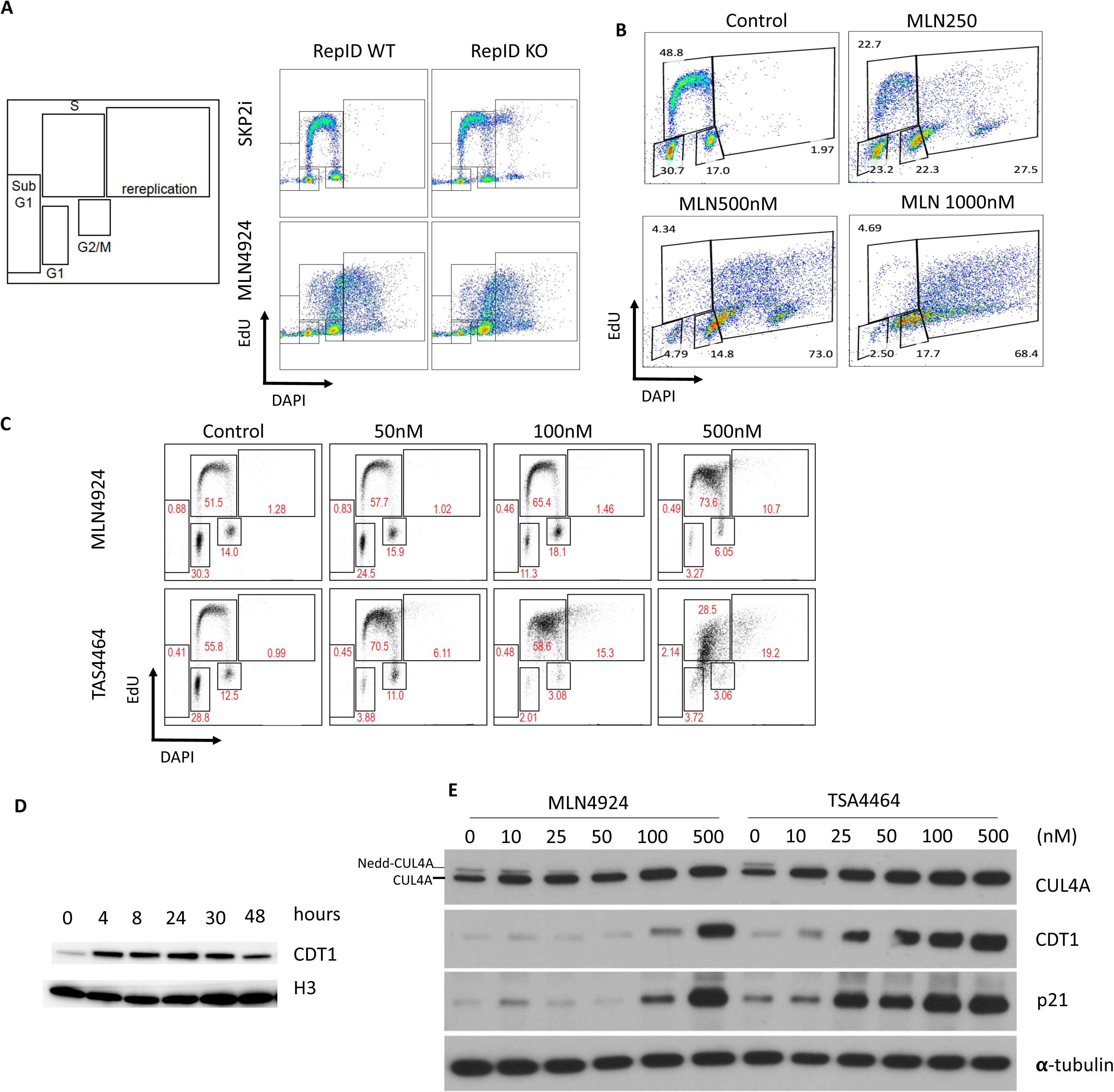
Different methods to induce massive re-replication in cancer cells. **a**, Shematic repsentation of gates used in the present study for cells at different cell cycle stages. More than G2/M cells are re-replication cells. Cell cycle profiles of RepID proficient and deficient U2OS cells treated with 50μM of SKP2 inhibitor for 3 days and 500nM of MLN4924 for 2 days. Cell cycle progression was monitored by flow cytometry. **b**, Cell cycle profiles of U2OS cells treated with the indicated concentration of MLN4924 for 48h. **c**, U2OS cells were treated with the indicated doses of MLN4924 or TAS4464 for 24h. Flow cytometry and immunoblotting were used to monitor cell cycle progression. **d,** HCT116 cells were treated with 250nM of MLN4924 for the indicated times and CDT1 levels were detected by immunoblotting. **e,** U2OS cells were treated with the indicated doses of MLN4924 or TAS4464 for 24h. Changes in CRL4 components and its substrates were detected by immunoblotting.

**Supplementary Fig 2.**
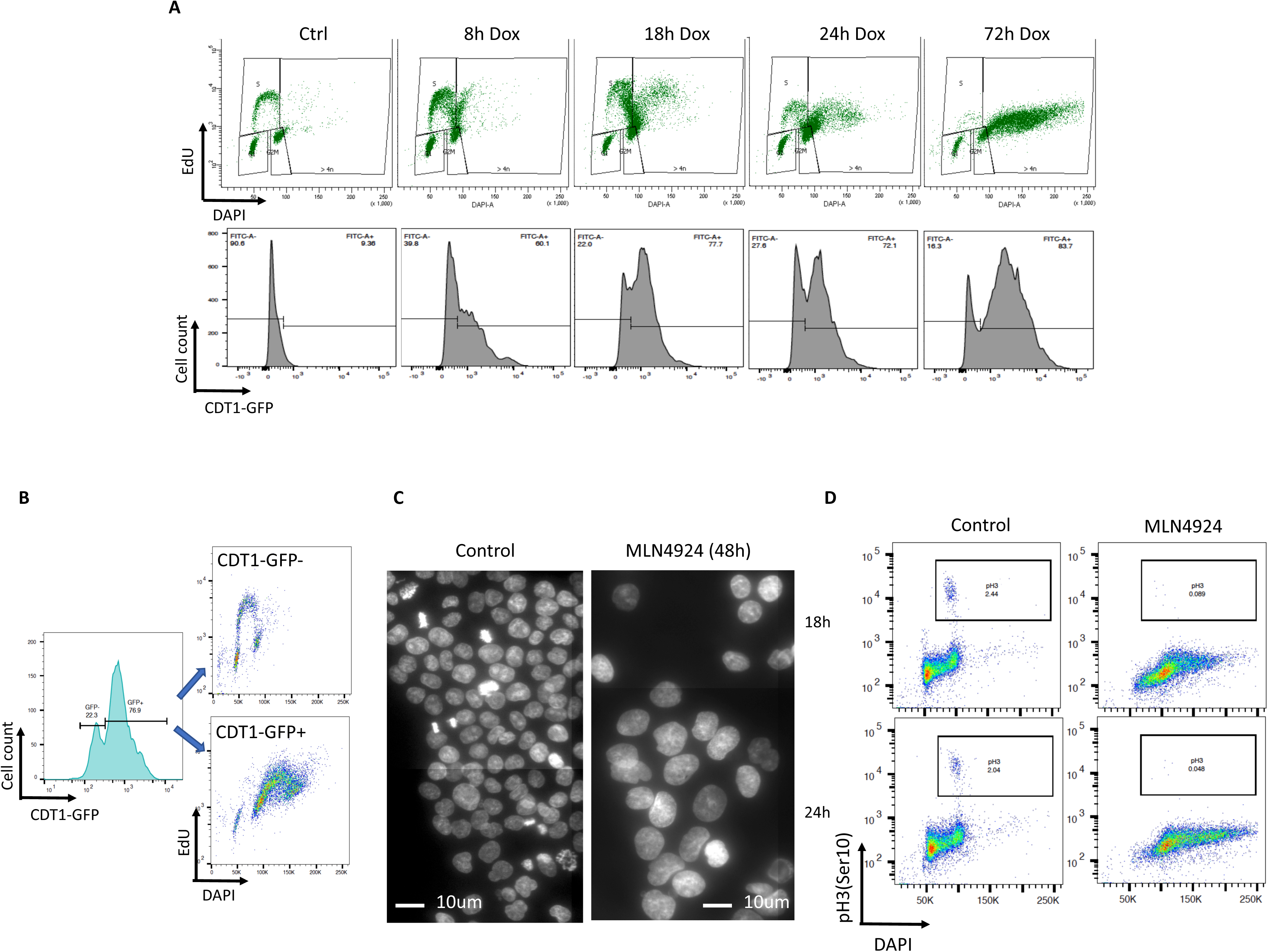
CDT1 stabilization induces DNA re-replication. **a**, Doxycycline inducible CDT1-GFP plasmid was stably transfected into U2OS cells (inducible CDT1-U2OS cells). Doxycycline 1ug/ml was added to induce CDT1 overexpression for the indicated times, and cell cycle progression (top) and CDT1-GFP level (bottom) were monitored by FACS. **b**, CDT1 OE was induced for 48h and the samples were gated based on CDT1-GFP levels (left). Cell cycle analyses of CDT1-GFP negative and positive cells (right) confirmed that cells that lost CDT1-GFP expression did not go re-replication. **c,** HCT116 cells were treated with 250nM of MLN4924 for 48h and stained by DAPI to visualize cells in mitosis. **d**, HCT116 cells were treated with 250nM of MLN4924 for 18 and 24h. Cells with phosphorylated H3 serine 10 (pH3(Ser10)) were detected by flow cytometry.

**Supplementary Fig 3.**
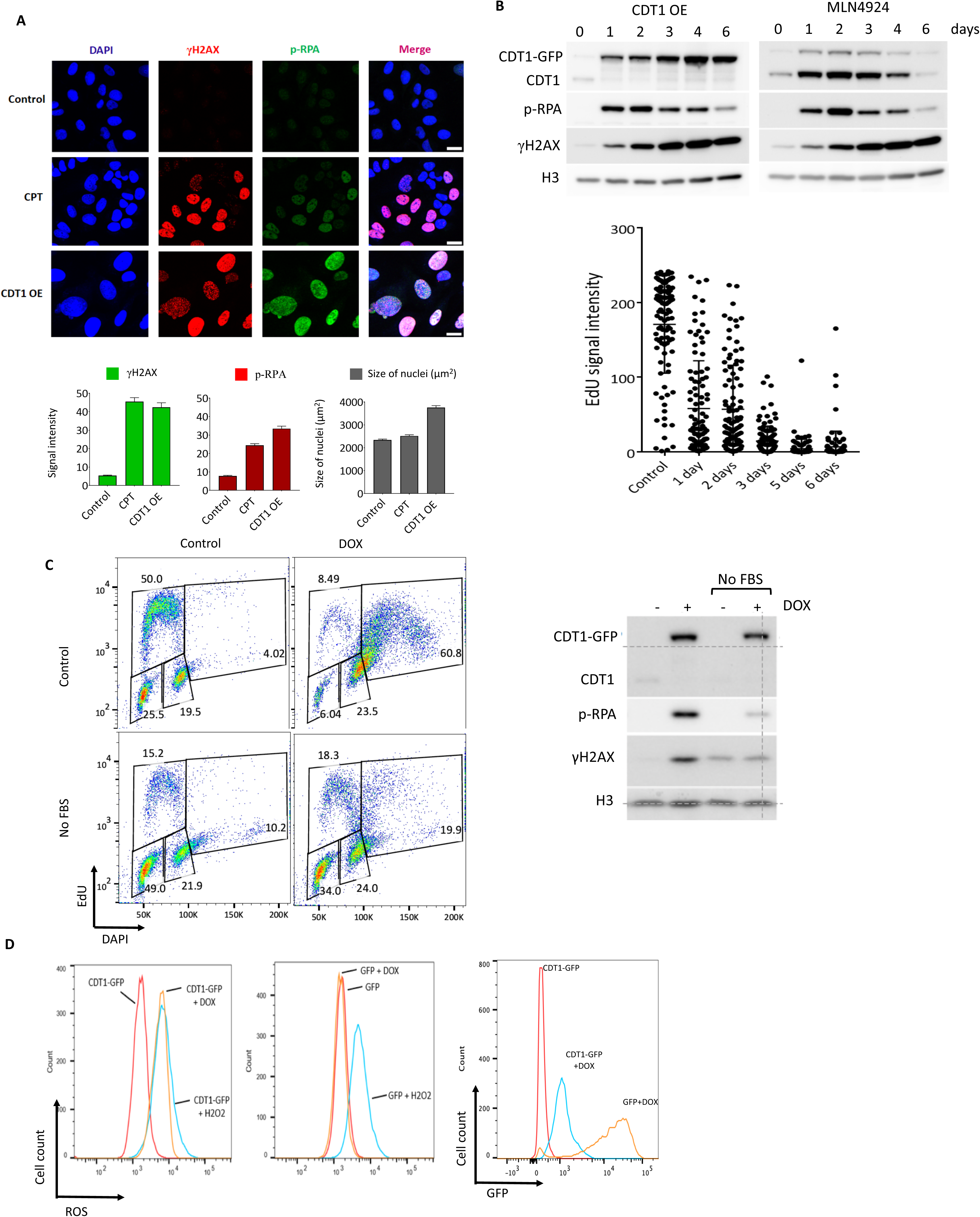
DNA re-replication induced by increased CDT1 level leads to replication stress and DNA damage. a, **a**, *top panel*, Representative images of inducible CDT1-U2OS cells treated with or without doxycycline for 48h, or with camptothecin (CPT) for 1 h. Cells were immunostained with γH2AX (red) and p-RPA (green). DNA was counterstained with DAPI. Scale bar: 20 µm. *Bottom panel*, Fluorescence intensities (arbitrary units) for p-RPA or γH2AX and nuclei size (µm^2^) are shown. b, Inducible CDT1-U2OS cells were treated with doxycycline (top left) or 500nM of MLN4924 (top right) for up to 6 days and changes in p-RPA, γH2AX and CDT1 levels were monitored by immunoblotting. Histone H3 was used as loading control. EdU signal intensities in re-replicating cells was monitored by immunostaining (bottom). CDT1 overexpression was induced in U2OS cells by doxycycline. EdU was added to cell cultures 45 min prior to cell collection and fixation at days 0, 1, 2, 3, 4 and 6 post-CDT1 induction. After EdU click reaction, cells were imaged by confocal microscopy. EdU intensities were measured in EdU positive cells using ImageJ (n>100 cells). c, Inducible CDT1-U2OS cells were cultured with and without fetal bovine serum (FBS) for 3 days prior to doxycycline treatment for 2 days. *Left panel*, Flow cytometry analysis to monitor cell cycle progression. *Right panel*, whole cell lysate analysis by Western to detect p-RPA, γH2AX and CDT1 levels. Histone H3 was used as loading control. d, Figure related to Fig 3d. Inducible GFP U2OS cells were treated with doxycycline for 48h to monitor the effect of GFP expression on ROS cellular production. ROS detection by flow cytometry in control cells (no DOX) and cells overexpressing CDT1-GFP (+DOX) (*left panel*) or GFP (+DOX) (*middle panel*). *Left panel*, detection of GFP levels in cells overexpressing CDT1-GFP or GFP (+DOX).

**Supplementary Fig. 4.**
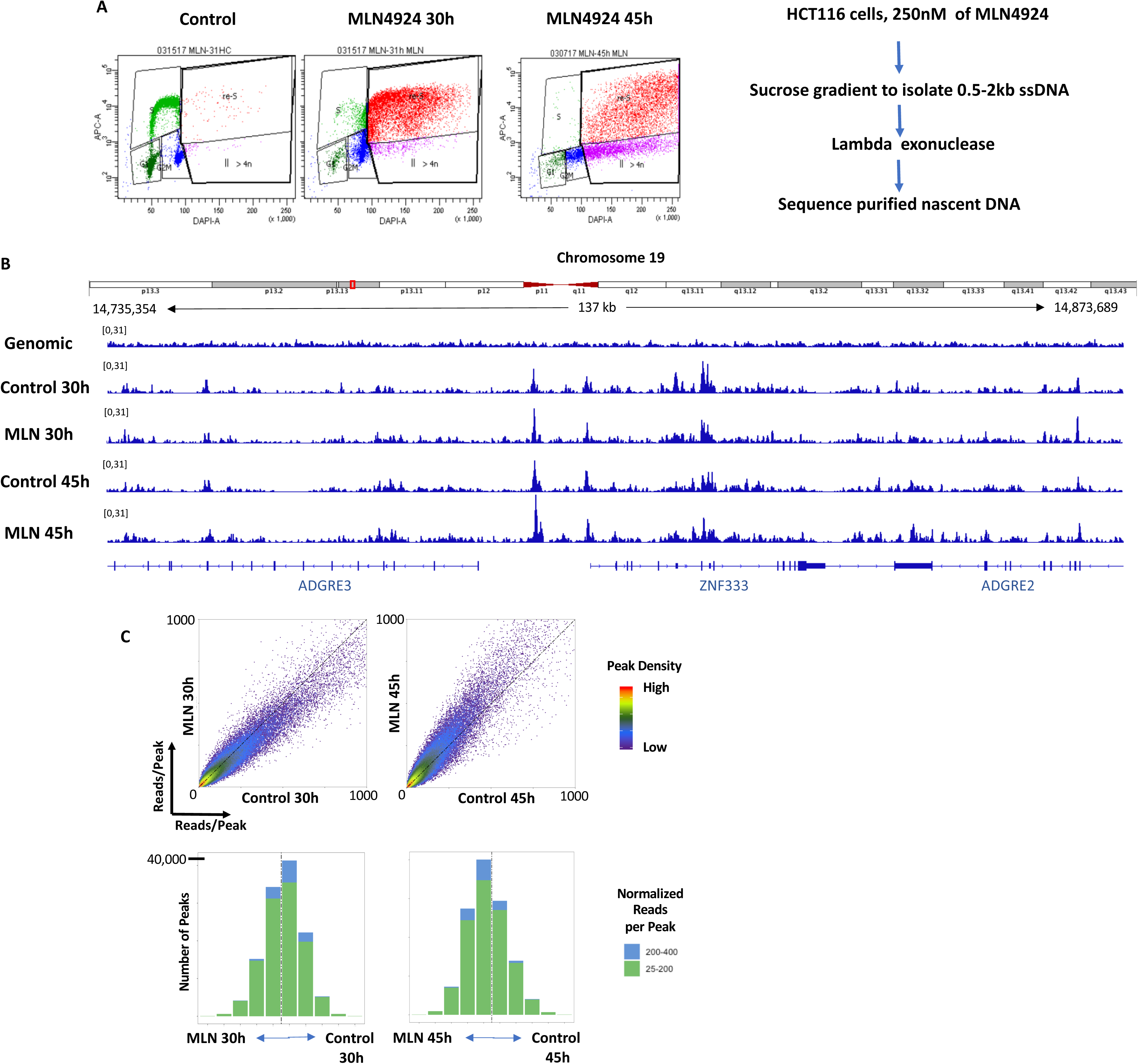
Re-replication is driven by the same set of origins utilized during normal mitotic growth. **a**, To map origins used during re-replication, HCT116 cells were treated with 250nM MLN4924 for 30h and 45h, timepoints for which almost all the replicating cells (30h) or all the replicating cells are in a re-replicating cycle (45h). Both control cells and MLN4924-treated cells were collected for nascent strand preparation, followed by next generation whole genome sequencing. Left: 2-dimensional flow cytometry to monitor cell cycle progression; right: experimental flow chart. **b**, An IGV screenshot shows representative nascent-Seq peaks, which represent origins, from both MLN4924-treated and untreated samples. Genomic: genomic DNA used as background to call replication origin peaks. **c**, Density plots of shared peaks between control cells and MLN4924-treated cells.

**Supplementary Fig 5.**
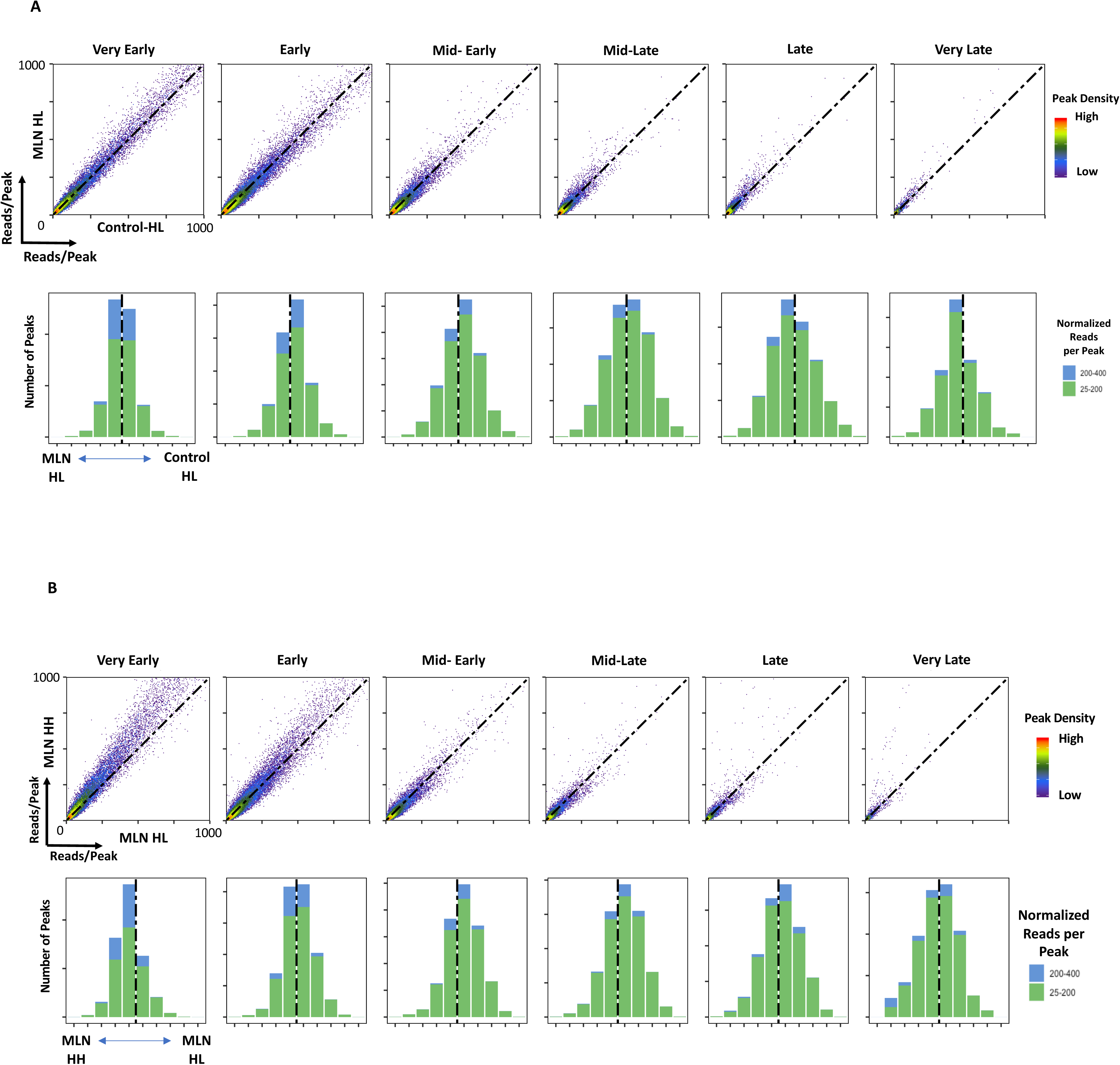
Re-replicating DNA is enriched at early genomic regions. Two sets of normal replication origins (MLN HL vs control HL) in a and one set of re-replication and one set of normal replication origins (MLN-HH vs MLN-HL) in b were compared as in Fig. 7d.

## References

1. Techer, H., Koundrioukoff, S., Nicolas, A. & Debatisse, M. The impact of replication stress on replication dynamics and DNA damage in vertebrate cells. Nat Rev Genet 18, 535–550 (2017).

2. Flach, J. et al. Replication stress is a potent driver of functional decline in ageing haematopoietic stem cells. Nature 512, 198–202 (2014).

3. Liontos, M. et al. Deregulated overexpression of hCdt1 and hCdc6 promotes malignant behavior. Cancer Res 67, 10899–10909 (2007).

4. Parker, M.W., Botchan, M.R. & Berger, J.M. Mechanisms and regulation of DNA replication initiation in eukaryotes. Crit Rev Biochem Mol Biol 52, 107–144 (2017).

5. Symeonidou, I.E. et al. Multi-step loading of human minichromosome maintenance proteins in live human cells. J Biol Chem 288, 35852–35867 (2013).

6. Parker, M.W. et al. A new class of disordered elements controls DNA replication through initiator self-assembly. Elife 8 (2019).

7. Ekundayo, B. & Bleichert, F. Origins of DNA replication. PLoS Genet 15, e1008320 (2019).

8. Utani, K. et al. Phosphorylated SIRT1 associates with replication origins to prevent excess replication initiation and preserve genomic stability. Nucleic Acids Res 45, 7807–7824 (2017).

9. Coleman, K.E. et al. Sequential replication-coupled destruction at G1/S ensures genome stability. Genes Dev 29, 1734–1746 (2015).

10. Walter, D. et al. SCF(Cyclin F)-dependent degradation of CDC6 suppresses DNA re-replication. Nat Commun 7, 10530 (2016).

11. Arias, E.E. & Walter, J.C. Replication-dependent destruction of Cdt1 limits DNA replication to a single round per cell cycle in Xenopus egg extracts. Genes Dev 19, 114–126 (2005).

12. Li, A. & Blow, J.J. Cdt1 downregulation by proteolysis and geminin inhibition prevents DNA re-replication in Xenopus. EMBO J 24, 395–404 (2005).

13. Smith, O.K. et al. Distinct epigenetic features of differentiation-regulated replication origins. Epigenetics Chromatin 9, 18 (2016).

14. Maiorano, D., Krasinska, L., Lutzmann, M. & Mechali, M. Recombinant Cdt1 induces rereplication of G2 nuclei in Xenopus egg extracts. Curr Biol 15, 146–153 (2005).

15. Pozo, P.N. & Cook, J.G. Regulation and Function of Cdt1; A Key Factor in Cell Proliferation and Genome Stability. Genes (Basel) 8 (2016).

16. Pozo, P.N. et al. Cdt1 variants reveal unanticipated aspects of interactions with Cyclin/CDK and MCM important for normal genome replication. Mol Biol Cell, mbcE18040242 (2018).

17. Nishitani, H. et al. Two E3 ubiquitin ligases, SCF-Skp2 and DDB1-Cul4, target human Cdt1 for proteolysis. EMBO J 25, 1126–1136 (2006).

18. Johansson, P. et al. SCF-FBXO31 E3 ligase targets DNA replication factor Cdt1 for proteolysis in the G2 phase of cell cycle to prevent re-replication. J Biol Chem 289, 18514–18525 (2014).

19. Havens, C.G. & Walter, J.C. Mechanism of CRL4(Cdt2), a PCNA-dependent E3 ubiquitin ligase. Genes Dev 25, 1568–1582 (2011).

20. Hayashi, A. et al. Direct binding of Cdt2 to PCNA is important for targeting the CRL4(Cdt2) E3 ligase activity to Cdt1. Life Sci Alliance 1, e201800238 (2018).

21. van Kempen, L.C. et al. The protein phosphatase 2A regulatory subunit PR70 is a gonosomal melanoma tumor suppressor gene. Sci Transl Med 8, 369ra177 (2016).

22. Jang, S.M., Redon, C.E. & Aladjem, M.I. Chromatin-Bound Cullin-Ring Ligases: Regulatory Roles in DNA Replication and Potential Targeting for Cancer Therapy. Front Mol Biosci 5, 19 (2018).

23. Munoz, S. et al. In Vivo DNA Re-replication Elicits Lethal Tissue Dysplasias. Cell Rep 19, 928–938 (2017).

24. Kerzendorfer, C. et al. Mutations in Cullin 4B result in a human syndrome associated with increased camptothecin-induced topoisomerase I-dependent DNA breaks. Hum Mol Genet 19, 1324–1334 (2010).

25. Zhang, Y. et al. A replicator-specific binding protein essential for site-specific initiation of DNA replication in mammalian cells. Nat Commun 7, 11748 (2016).

26. Jang, S.M. et al. The replication initiation determinant protein (RepID) modulates replication by recruiting CUL4 to chromatin. Nat Commun 9, 2782 (2018).

27. Fu, H. et al. Methylation of histone H3 on lysine 79 associates with a group of replication origins and helps limit DNA replication once per cell cycle. PLoS Genet 9, e1003542 (2013).

28. Lin, J.J., Milhollen, M.A., Smith, P.G., Narayanan, U. & Dutta, A. NEDD8-targeting drug MLN4924 elicits DNA rereplication by stabilizing Cdt1 in S phase, triggering checkpoint activation, apoptosis, and senescence in cancer cells. Cancer Res 70, 10310–10320 (2010).

29. Petrakis, T.G. et al. Exploring and exploiting the systemic effects of deregulated replication licensing. Semin Cancer Biol 37-38, 3-15 (2016).

30. Best, S. et al. Targeting ubiquitin-activating enzyme induces ER stress-mediated apoptosis in B-cell lymphoma cells. Blood Adv 3, 51–62 (2019).

31. Vanderdys, V. et al. The Neddylation Inhibitor Pevonedistat (MLN4924) Suppresses and Radiosensitizes Head and Neck Squamous Carcinoma Cells and Tumors. Mol Cancer Ther 17, 368–380 (2018).

32. Blank, J.L. et al. Novel DNA damage checkpoints mediating cell death induced by the NEDD8-activating enzyme inhibitor MLN4924. Cancer Res 73, 225–234 (2013).

33. Yoshimura, C., et al. TAS4464, A Highly Potent and Selective Inhibitor of NEDD8-Activating Enzyme, Suppresses Neddylation and Shows Antitumor Activity in Diverse Cancer Models. Mol Cancer Ther 18, 1205–1216 (2019).

34. Varma, D. et al. Recruitment of the human Cdt1 replication licensing protein by the loop domain of Hec1 is required for stable kinetochore-microtubule attachment. Nat Cell Biol 14, 593–603 (2012).

35. Fu, H. et al. The DNA repair endonuclease Mus81 facilitates fast DNA replication in the absence of exogenous damage. Nat Commun 6, 6746 (2015).

36. Gong, D. & Ferrell, J.E., Jr. The roles of cyclin A2, B1, and B2 in early and late mitotic events. Mol Biol Cell 21, 3149–3161 (2010).

37. Hendzel, M.J. et al. Mitosis-specific phosphorylation of histone H3 initiates primarily within pericentromeric heterochromatin during G2 and spreads in an ordered fashion coincident with mitotic chromosome condensation. Chromosoma 106, 348–360 (1997).

38. Meselson, M. & Stahl, F.W. The Replication of DNA in Escherichia Coli. Proc Natl Acad Sci U S A 44, 671–682 (1958).

39. Fu, H. et al. Mapping replication origin sequences in eukaryotic chromosomes. Curr Protoc Cell Biol 65, 22 20 21–17 (2014).

40. Martin, M.M. et al. Genome-wide depletion of replication initiation events in highly transcribed regions. Genome Res 21, 1822–1832 (2011).

41. Koren, A. et al. Genetic variation in human DNA replication timing. Cell 159, 1015–1026 (2014).

42. Mukhopadhyay, R. et al. Allele-specific genome-wide profiling in human primary erythroblasts reveal replication program organization. PLoS Genet 10, e1004319 (2014).

43. Ullah, Z., Lee, C.Y., Lilly, M.A. & DePamphilis, M.L. Developmentally programmed endoreduplication in animals. Cell Cycle 8, 1501–1509 (2009).

44. Vaziri, C. et al. A p53-dependent checkpoint pathway prevents rereplication. Mol Cell 11, 997–1008 (2003).

45. Abbas, T., Keaton, M.A. & Dutta, A. Genomic instability in cancer. Cold Spring Harb Perspect Biol 5, a012914 (2013).

46. Abbas, T. & Dutta, A. Regulation of Mammalian DNA Replication via the Ubiquitin-Proteasome System. Adv Exp Med Biol 1042, 421–454 (2017).

47. Zhu, W. & Depamphilis, M.L. Selective killing of cancer cells by suppression of geminin activity. Cancer Res 69, 4870–4877 (2009).

48. Anglana, M., Apiou, F., Bensimon, A. & Debatisse, M. Dynamics of DNA replication in mammalian somatic cells: nucleotide pool modulates origin choice and interorigin spacing. Cell 114, 385–394 (2003).

49. McIntosh, D. & Blow, J.J. Dormant origins, the licensing checkpoint, and the response to replicative stresses. Cold Spring Harb Perspect Biol 4 (2012).

50. Alver, R.C., Chadha, G.S. & Blow, J.J. The contribution of dormant origins to genome stability: from cell biology to human genetics. DNA Repair (Amst) 19, 182–189 (2014).

51. Shima, N. & Pederson, K.D. Dormant origins as a built-in safeguard in eukaryotic DNA replication against genome instability and disease development. DNA Repair (Amst) 56, 166–173 (2017).

52. Moiseeva, T.N. et al. An ATR and CHK1 kinase signaling mechanism that limits origin firing during unperturbed DNA replication. Proc Natl Acad Sci U S A 116, 13374–13383 (2019).

53. Aymard, F. et al. Genome-wide mapping of long-range contacts unveils clustering of DNA double-strand breaks at damaged active genes. Nat Struct Mol Biol 24, 353–361 (2017).

54. Clouaire, T. et al. Comprehensive Mapping of Histone Modifications at DNA Double-Strand Breaks Deciphers Repair Pathway Chromatin Signatures. Mol Cell 72, 250–262 e256 (2018).

55. Irony-Tur Sinai, M. & Kerem, B. Genomic instability in fragile sites-still adding the pieces. Genes Chromosomes Cancer 58, 295–304 (2019).

56. Alvarez, S. et al. Replication stress caused by low MCM expression limits fetal erythropoiesis and hematopoietic stem cell functionality. Nat Commun 6, 8548 (2015).

57. Bolger, A.M., Lohse, M. & Usadel, B. Trimmomatic: a flexible trimmer for Illumina sequence data. Bioinformatics 30, 2114–2120 (2014).

58. Li, H. & Durbin, R. Fast and accurate short read alignment with Burrows-Wheeler transform. Bioinformatics 25, 1754–1760 (2009).

59. Zhang, Y. et al. Model-based analysis of ChIP-Seq (MACS). Genome Biol 9, R137 (2008).

